# Barley yield formation under abiotic stress depends on the interplay between flowering time genes and environmental cues

**DOI:** 10.1101/488080

**Authors:** Mathias Wiegmann, Andreas Maurer, Anh Pham, Timothy J. March, Ayed Al-Abdallat, William T.B. Thomas, Hazel J. Bull, Mohammed Shahid, Jason Eglinton, Michael Baum, Andrew J. Flavell, Mark Tester, Klaus Pillen

## Abstract

Since the dawn of agriculture, crop yield has always been impaired through abiotic stresses. In a field trial across five locations worldwide, we tested three abiotic stresses, nitrogen deficiency, drought and salinity, using HEB-YIELD, a selected subset of the wild barley nested association mapping population HEB-25. We show that barley flowering time genes *Ppd-H1, Sdw1, Vrn-H1* and *Vrn-H3* exert pleiotropic effects on plant development and grain yield. Under field conditions, these effects are strongly influenced by environmental cues like day length and temperature. For example, in Al-Karak, Jordan, the day length-sensitive wild barley allele of *Ppd-H1* was associated with an increase of grain yield by up to 30% compared to the insensitive elite barley allele. The observed yield increase is accompanied by pleiotropic effects of *Ppd-H1* resulting in shorter life cycle, extended grain filling period and increased grain size. Our study indicates that the adequate timing of plant development is crucial to maximize yield formation under harsh environmental conditions. We provide evidence that wild barley germplasm, introgressed into elite barley cultivars, can be utilized to improve grain yield. The presented knowledge may be transferred to related crop species like wheat and rice securing the rising global food demand for cereals.

## Introduction

One of the major challenges that mankind faces is the ability to feed the ever-growing population, especially in the face of increased stresses due to climate change and reduced availability of arable land^1,2^. Different climate prediction models indicate severe effects for large parts of Africa, the Arabian Peninsula and Central South America^3,4^, where barley (*Hordeum vulgare* ssp. *vulgare*) still has an crucial role as human food^5^. Barley is mainly used for animal feed and for malt production in large parts of the world. It represents the fourth most important cereal crop on a global scale^5,6^.

Barley inherently exhibits a higher level of abiotic stress tolerance than other crops^7–9^, which offers the possibility to extend its future production to areas suffering from climate change. Furthermore, the relatively simple diploid genetics of barley and the tight relationship between the members of the *Triticeae* tribe facilitate the transfer of knowledge gained from barley research to other major cereals, for instance, bread wheat, durum wheat and rye^10^. Wild barley (*Hordeum vulgare* ssp. *spontaneum*), originating from the Fertile Crescent and from a second area some 1,500–3,000 km farther east, was used to domesticate modern elite barley (*Hordeum vulgare* ssp. *vulgare*) more than 10,000 years ago^11–13^. The usefulness of wild germplasm for future breeding has often been emphasized^14–16^, mostly as a source to improve biotic resistance and abiotic stress tolerance rather than to directly increase grain yield^17^. Recent studies in wild barley indicate the existence of vast phenological variation for important agronomic traits^18–25^. Wild barely may thus be an appropriate source to replenish the barley gene pool with novel genetic variation. This variation may be valuable to cope with the challenges arising from climate change^26^.

Grain yield depends on developmental phases of a plant’s life cycle^27^. In this regard, flowering time is a key event as plants shift from vegetative to reproductive growth, moving towards providing the harvestable yield^28–30^. The optimal timing of this event is crucial as it should occur in the absence of adverse effects like abiotic stresses^31^ but also ensuring completion of yield accumulation without encountering further adverse effects in most growing seasons. Therefore, the targeted timing of this phase provides one approach to improve stress tolerance, through stress avoidance, and thus to increase grain yield^32^. Flowering time is mainly controlled by environmental cues like day length (photoperiod) and temperature (especially the exposure to cold temperatures, also termed vernalization)^33–35^. Flowering time is highly heritable and, so far, several major genes controlling flowering time have been discovered in model species and in crop plants^36^. Generally, flowering time genes are classified into at least three families: [I] photoperiod genes (e.g. *Ppd-H1*) ^37^, [II] vernalization genes (e.g. *Vrn-H1* and *Vrn-H3*)^33,38^ and [III] earliness *per se* (*eps*) genes, the last controlling flowering independently from photoperiod and temperature (e.g. *Sdw1*)^39,40^.

Here, we present data of a large field study with the HEB-YIELD population, a selected subset of the wild barley nested association mapping (NAM) population HEB-25^18^. The aim of the study was to examine the interplay between flowering time, stress tolerance and yield. For this purpose, HEB-YIELD was studied at five locations worldwide and during two years under locally relevant abiotic stress conditions. We investigated the role of known flowering time genes on developmental and yield-related traits, as well as how they account for yield and stress tolerance.

## Results and discussion

### HEB-YIELD exhibits strong phenotypic variation as well as environmental and treatment variation

The wild barley introgression population HEB-YIELD comprises a diverse subset of lines selected from the NAM population HEB-25^18^ (Supplementary Table S1). We studied eleven agronomically traits in a HEB-YIELD trial conducted in Dundee, Halle, Al-Karak, Dubai and Adelaide (Figure 1; Supplementary Table S2a), where climate data for day length, temperature and precipitation varied considerably between locations Supplementary Figure S1 and S2). The parameters studied included developmental and yield-related traits, used to capture growth variation among HEB-YIELD lines (Supplementary Table S3). At each location the traits were measured under site-specific abiotic stress conditions, i.e. nitrogen deficiency in Dundee and Halle, drought stress in Al-Karak and Adelaide and salt stress in Dubai. In total, 3,207 field plots were evaluated over all sites, seasons, treatments, and replicates (Supplementary Table S2c and S4a). Considerable phenotypic variation within locations and treatments was observed for all investigated traits (Figure 2; Supplementary Table S5).

**Figure 1.**
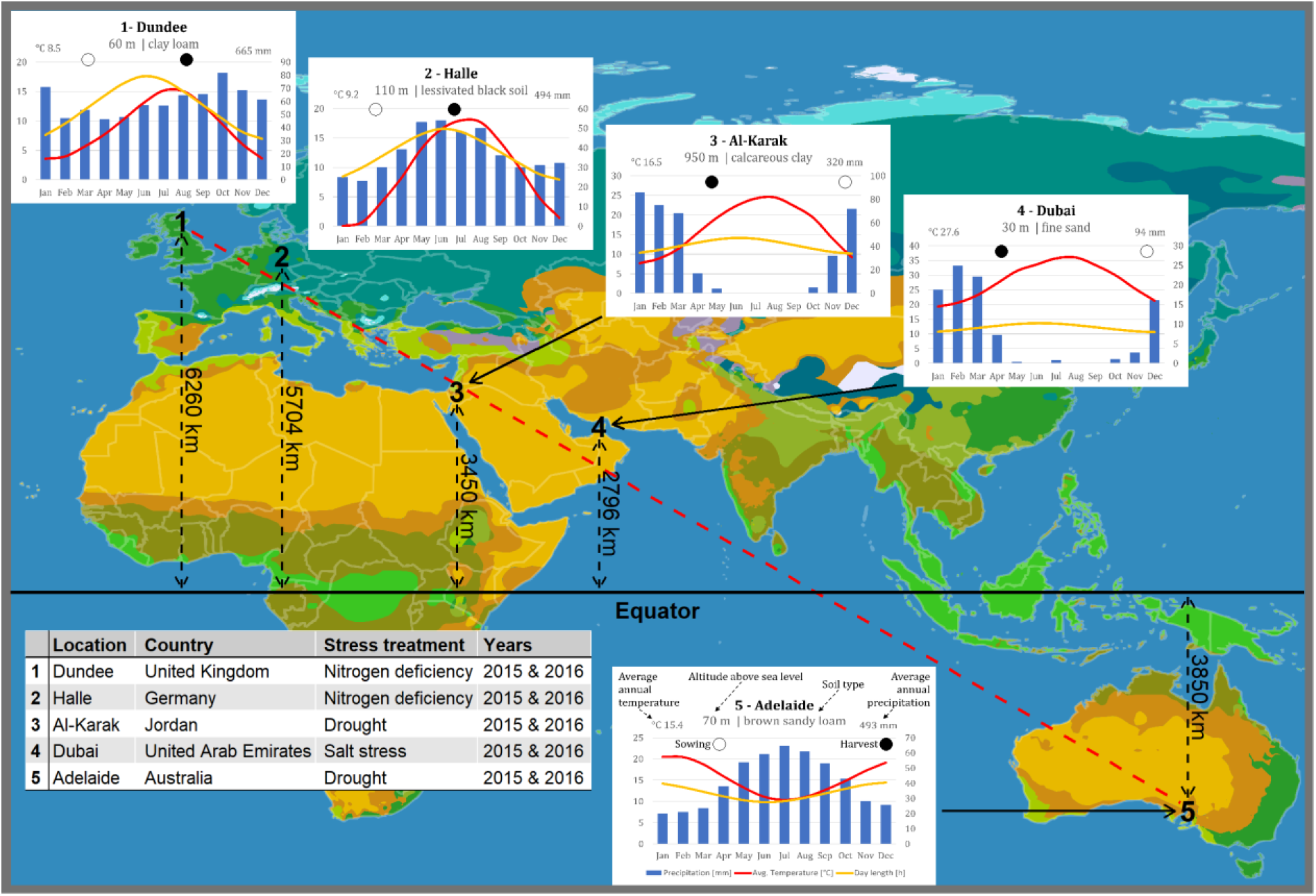
Global macroclimate map with information on the five experimental locations. The position of the five (1-5) test locations are indicated on a simplified map of the Köppen-Geiger climate classification system. General information about the test locations are given in the table on the lower left-hand side including the nearest town, country, stress treatment and the years of field trials. Insets next to map positions depicts long-term climate information for each test location. The average monthly precipitation in millimeters (blue bars), the average monthly temperature in degrees Celsius (red line) and the course of the day length during the year in hours (yellow line) are displayed. In addition, the sowing and harvesting dates are indicated with empty and filled circles, respectively. The Adelaide inset on the right-hand side serves as a legend for the insets. Map source: http://en.wikipedia.org/wiki/World_map.

**Figure 2.**
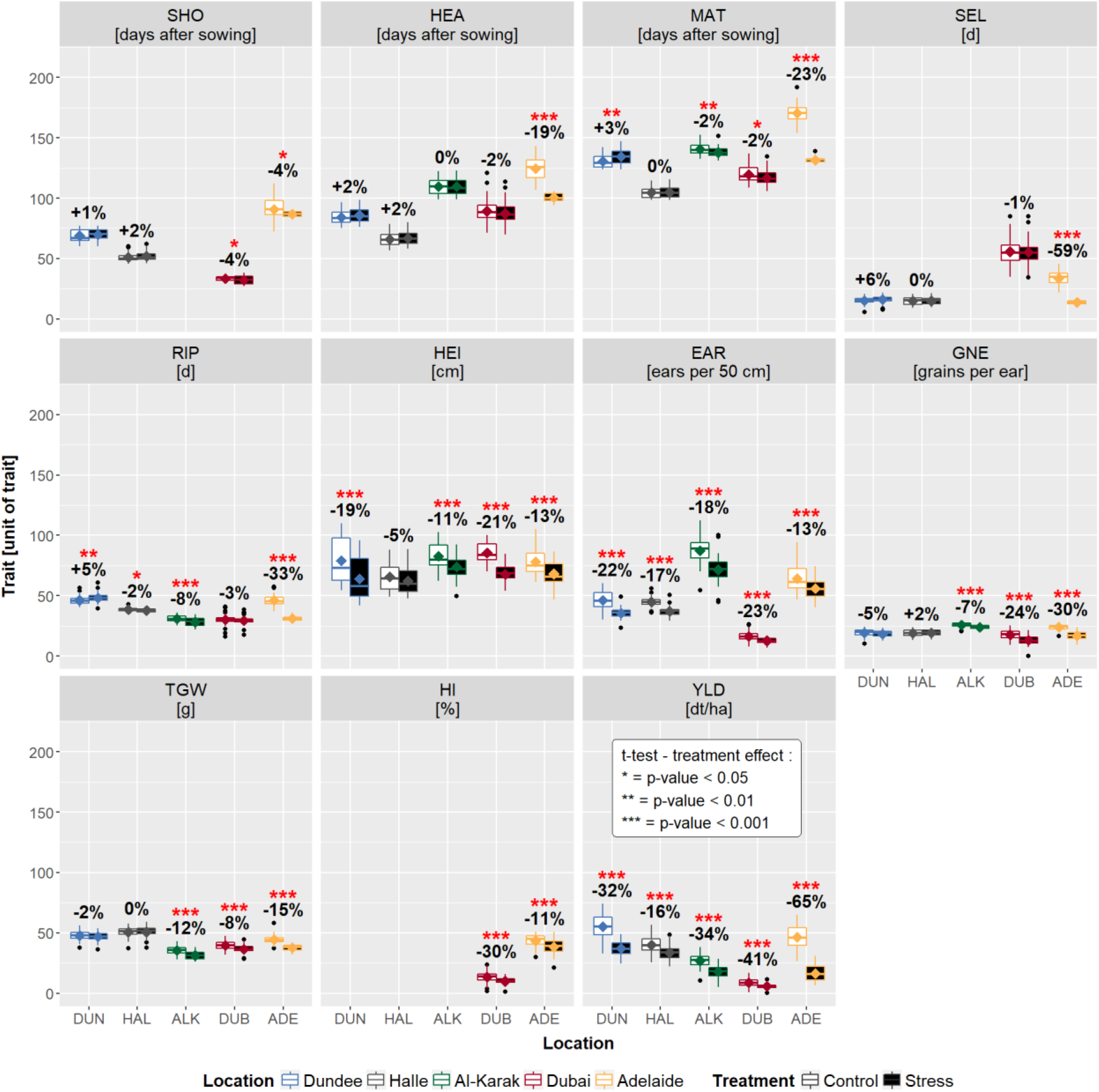
Box-Whisker plots illustrating HEB-YIELD trait variation per location and treatment. Trait names and trait units are indicated in the grey rectangle above each subplot. Trait abbreviations are listed in Supplementary Table S3. The locations Dundee (DUN), Halle (HAL), Al-Karak (ALK), Dubai (DUB) and Adelaide (ADE) are indicated with blue, grey, green, red and yellow box-whiskers, respectively, and, in addition, at the bottom of the plot. Empty and filled boxes refer to control and stress treatments, respectively. Significant differences between treatments are indicated with red asterisks above boxes with *: *P* < 0.05, **: *P* < 0.01 and ***: *P* < 0.001. The relative increase/decrease (in %) of the stress treatment compared to the control treatment is given below the asterisks.

The ANOVA revealed that all investigated factors (genotype, year and location) were significant for all traits except plant height (HEI) where the year effect was not significant (Supplementary Table S6a). Interestingly, only the traits shoot elongation phase (SEL) and HEI showed comparable values across locations, whereas for the majority of traits pronounced location effects were observed (Supplementary Table S6c). For instance, flowering time varied from 57 to 144 days and grain yield from 0.14 dt/ha to 74 dt/ha (Supplementary Table S5), reflecting a strong diversity in yield potential among the trial sites (Figure 2).

Irrespective of the diverging agricultural practices at the trial sites, developmental trait heritabilities were high with an average of 0.87, ranging from 0.10 (ripening phase (RIP) under control treatment in Al-Karak) to 0.99 (shooting (SHO) under control treatment in Adelaide as well as flowering (HEA) and SEL under both treatments in Dubai, Supplementary Table S5). In general, yield-related traits revealed lower heritabilities with an average of 0.65. The most complex trait, grain yield (YLD), revealed average heritabilities of 0.73, ranging from 0.05 (YLD under stress treatment in Dubai) to 0.93 (YLD under control treatment in Dundee).

### Trait performance in HEB-YIELD is usually a linear transformation from control to stress treatments indicating low genotype by treatment interaction

To gain insights into how abiotic stresses may affect plant development and grain yield, we cultivated HEB-YIELD under contrasting stress conditions, which are relevant for the respective test locations (Supplementary Table S2c). The applied stresses exhibited only minor effects on plant development traits except for HEI. In contrast, strong effects on all measured yield-related traits were observed at all test locations, for instance, reducing yield under stress between 16 % in Halle and 65 % in Adelaide (Figure 2; Supplementary Table S5). We observed a weak trend, that HEB-YIELD lines under drought and salt stress exhibited an accelerated plant development, presumably to escape the stress condition, which is in agreement with other studies in cereals^32,45^. Based on our findings we suggest that plant development in the wild barley population HEB-YIELD is mainly determined by genetic factors and to a lesser extend modified by abiotic stresses. This is further supported by the observation that plant developmental traits showed a nearly linear shift between control and stress conditions, as indicated by high correlation coefficients (0.99 > r > 0.59) between stress and control treatments of developmental traits, except for SEL in Adelaide (r=0.12; Table 1; Supplementary Table S7a).

**Table 1.**
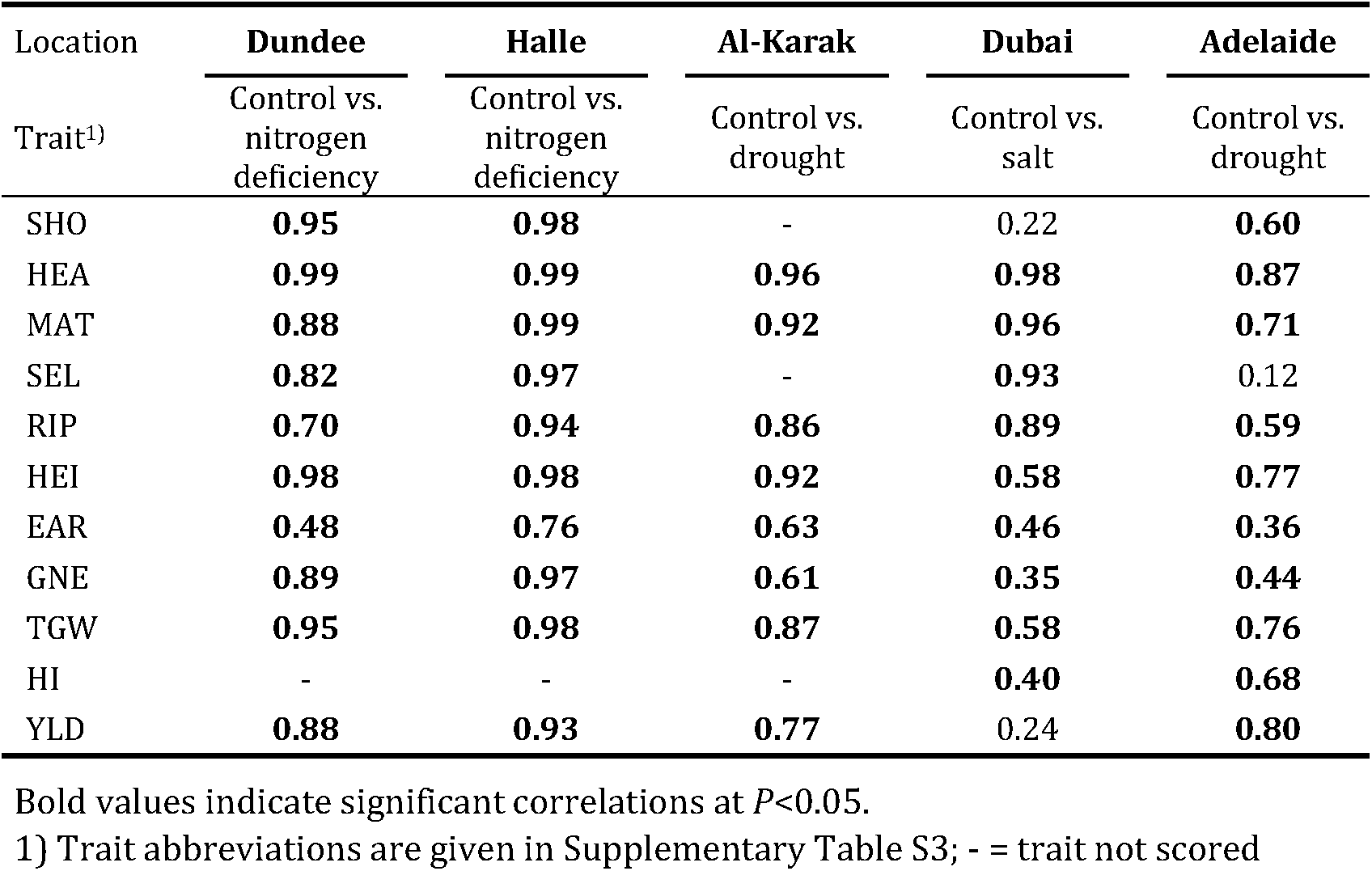
Location-specific Pearson correlation coefficients (*r*) within trait, measured under control versus stress condition

| Location | Dundee | Halle | Al-Karak | Dubai | Adelaide |
| --- | --- | --- | --- | --- | --- |
| Trait<sup>1)</sup> | Control vs.<br>nitrogen<br>deficiency | Control vs.<br>nitrogen<br>deficiency | Control vs.<br>drought | Control vs.<br>salt | Control vs.<br>drought |
| SHO | **0.95** | **0.98** | - | 0.22 | **0.60** |
| HEA | **0.99** | **0.99** | **0.96** | **0.98** | **0.87** |
| MAT | **0.88** | **0.99** | **0.92** | **0.96** | **0.71** |
| SEL | **0.82** | **0.97** | - | **0.93** | 0.12 |
| RIP | **0.70** | **0.94** | **0.86** | **0.89** | **0.59** |
| HEI | **0.98** | **0.98** | **0.92** | **0.58** | **0.77** |
| EAR | **0.48** | **0.76** | **0.63** | **0.46** | **0.36** |
| GNE | **0.89** | **0.97** | **0.61** | **0.35** | **0.44** |
| TGW | **0.95** | **0.98** | **0.87** | **0.58** | **0.76** |
| HI | - | - | - | **0.40** | **0.68** |
| YLD | **0.88** | **0.93** | **0.77** | 0.24 | **0.80** |

Bold values indicate significant correlations at $P < 0.05$ .
1) Trait abbreviations are given in Supplementary Table S3; - = trait not scored

### Grain yield correlations indicate that yield formation depends on a location-specific interplay between developmental traits and yield components

We observed Pearson correlations coefficients between plant developmental stages shooting, flowering and maturity ranging from r=0.67 to r=0.96 (apart from shooting correlations in Dubai Supplementary Tables S6b and S7b), indicating a nearly colinear regulation of plant developmental phases. Thus, HEB-YIELD lines early or late in shooting have the tendency to stay early or late respectively until maturity. This observation is in agreement with previous findings in the wild barley NAM population HEB-25, studied in Halle^20^ and Dundee^25^. Consequently, early developmental stages may be used as an indirect criterion to select HEB-YIELD lines for early or late maturity.

Following these findings, we explored the relationship between plant development and yield formation in HEB-YIELD (Table 2). We observed a trend that late plant development is beneficial for increased grain yield under Dundee, Halle and Adelaide growth conditions, indicated by positive correlation coefficients of r(HEAxYLD) = 0.59/0.66, 0.32/0.20 and 0.57/0.51, respectively, under control/stress treatments. This trend fits the general observation that late lines have the potential to exploit a prolonged growing season if the environmental conditions including temperature and precipitation are beneficial^29,46,47^. In contrast, under the harsh environmental conditions at Al-Karak and Dubai, HEB-YIELD lines with accelerated plant development were favored. Consequently, we observed negative correlations between flowering and grain yield at Al-Karak and Dubai with r(HEAxYLD) = −0.30/−0.72 and −0.51/−0.44, respectively, under control/stress treatments (Table 2). Here, elevated temperatures and low rainfall restricted plant growth to a few months and thus earliness is a major breeding goal^48,49^. In future, this situation may intensify, since climate change is expected to further shorten the growing period in drought and heat prone locations like in Jordan^26,32,50^.

**Table 2.**
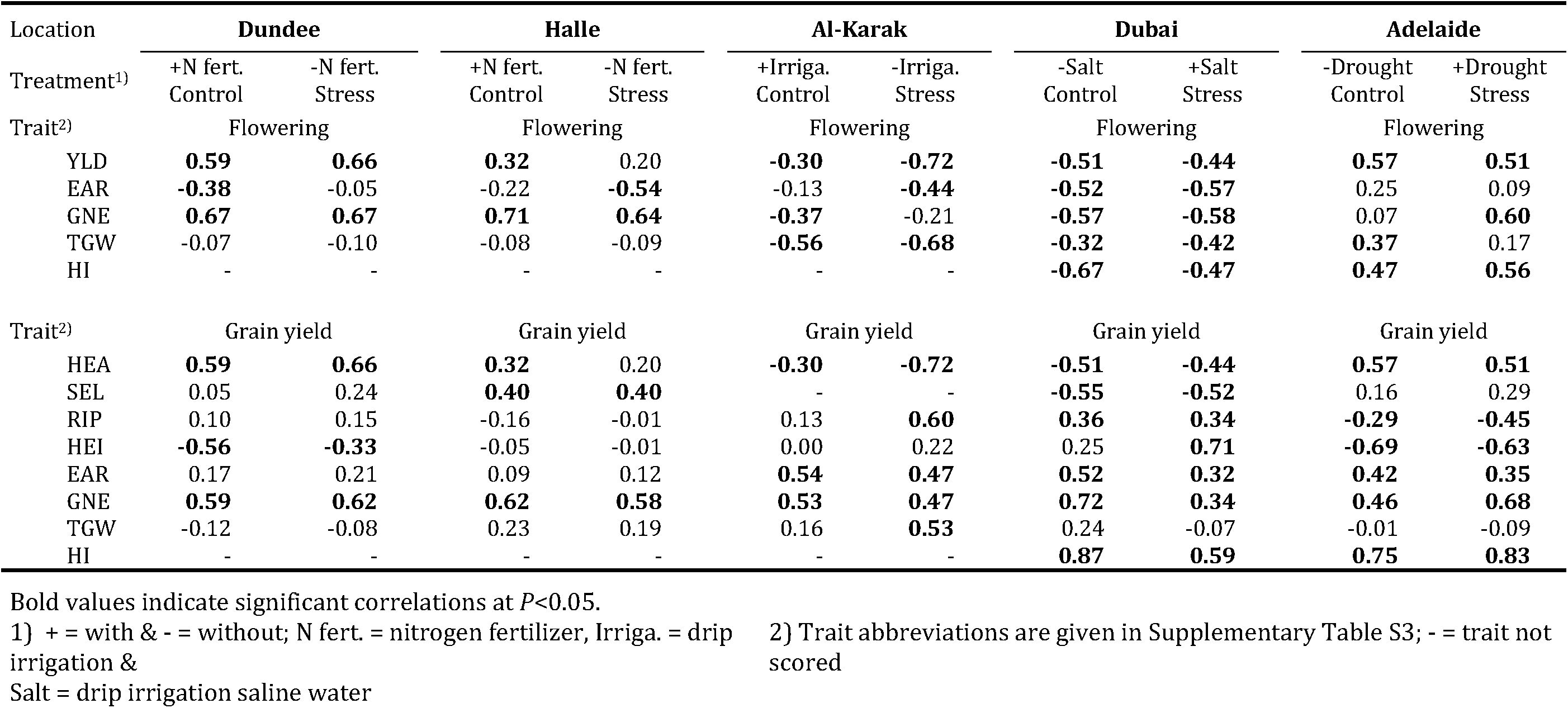
Location and treatment specific Pearson correlation coefficients (*r*) between plant developmental traits and flowering time (upper part) and grain yield (lower part), respectively

We also observed strong location-specific correlations between flowering time and yield components. For example, in Halle and Dundee, flowering time was positively correlated with grain number per ear (GNE), with r(HEAxGNE) = 0.67/0.67 and 0.71/0.64, respectively (Table 2). Here the extended vegetative growth phase allowed more spikelet primordia to be maintained. In contrast, in Al-Karak flowering time negatively affected thousand grain weight (TGW) with r(HEAxTGW) = −0.56/−0.68, reflecting a grain filling penalty for later flowering genotypes. These findings suggest that flowering time controls final grain yield to a certain degree. By comparing correlations between grain yield and yield components, we observed that apparently GNE is the key determinant of grain yield in HEB-YIELD – irrespective of location and treatment. This result is in agreement with earlier studies^51,52^, indicating that any increase in number of grains may also improve grain yield^53,54^. We thus reason that improving GNE may offer the best route to increase grain yield in HEB-YIELD independent of the environmental conditions.

The highest positive correlations of yield were found with harvest index (HI; scored only in Dubai and Adelaide with r(YLDxHI) = 0.87 and 0.83, respectively, Table 2). A previous study noted the importance of increasing harvest index to improve yield during the past century^55^. However, a further improvement of grain yield through raising harvest index may be a dead end, since barley is supposed to have reached an optimum with a harvest index of approximately 0.62^55,56^. Therefore, future grain yield improvements may be achieved through increasing plant biomass^55,57^. This suggestion is in accordance with our finding that grain yield exhibited a slightly positive correlation with shoot elongation phase in those environments where lateness was beneficial to increase yield (Table 2). During shoot elongation, which captures the growth period between establishing awn primordia and ear emergence, the leaf growth rate and the potential grain number per area are defined^51,53,58^. An extended shoot elongation phase may thus improve grain yield by increasing leaf size, i.e. biomass, and grain number per area. On the other hand, ripening phase under drought stress exhibited positive and negative correlations with grain yield in Al-Karak and in Adelaide, respectively. Whereas the Adelaide finding fits the assumption that early maturity and thus a short ripening phase may improve grain yield under terminal drought, the Al-Karak finding is unexpected. Under drought stress conditions in Al-Karak an extended ripening phase was associated with an increase in grain weight, ultimately resulting in elevated grain yields. We conclude that fine-tuning of plant development, especially their sub-phases, may contribute to a better adaptation of improved varieties to their target environment. The latter notion is supported by the finding that in the first instance climate change is expected to impair flowering time^32^, which is crucial for plant adaptation and yield formation^53,59^. In addition, our stress treatments confirmed the known association between inflorescence development and stress tolerance/avoidance^32,60^. This offers the possibility to use the genetically relatively well-understood trait flowering time as a proxy to select for improved grain yield under abiotic stresses^61^.

### Flowering time genes exhibit pleiotropic effects on yield formation in HEB-YIELD

In order to explore the interplay between flowering time regulation and yield formation, we investigated the effects of four major flowering time genes, *Ppd-H1, Sdw1, Vrn-H1* and *Vrn-H3,* in HEB-YIELD (Supplementary Table S8a). The relevance of these candidate genes has been reported in various studies^34,36^, including the wild barley NAM population HEB-25^18,20,22,25^. The wild barley lines of HEB-YIELD were selected to compare the effects of wild and cultivated alleles at these four flowering time loci (Supplementary Table S1). In the following, we report on the pleiotropic effects associated with the four flowering genes studied.

#### Ppd-H1

Flowering under long days is promoted by the photoperiod responsive *Ppd-H1* (*PHOTOPERIOD-H1*) allele, an orthologue of the Arabidopsis pseudoresponse regulator gene *PRR7*, which is present in wild barley and winter barley cultivars^37^. In contrast, spring barley cultivars like Barke possess the recessive non-responsive *ppd-H1* allele, resulting in late flowering. During early plant development, the photoperiod signal is transmitted from the circadian clock oscillator *Ppd-H1* through mediation of the CONSTANS (CO) protein to the floral inducer *Vrn-H3,* an orthologue of the Arabidopsis *FLOWERING LOCUS T* (*FT*) gene.

Among the candidate genes, *Ppd-H1* revealed the most pronounced effects on plant development in Dundee, Halle and Al-Karak (Figure 3; Supplementary Table S8a). This finding is in accordance with several other studies conducted in barley^20,23,25^. At these locations, the wild allele of *Ppd-H1* accelerated plant development in HEB-YIELD (SHO, HEA and maturity (MAT)) with a maximum effect of −9.0 days in Halle. In contrast, no significant effect of the *Ppd-H1* wild allele was observed in Dubai and Adelaide. Most wild barley accessions carry the dominant allele, which is responsive to a long day photoperiod, accelerating plant development through upregulating of *Vrn-H3*/*HvFT1*^62,63^. One possible explanation for contrasting effects between locations is the different day lengths at these sites. Dundee and Halle are more than 5,700 km distant from the equator and are clearly exposed to long day conditions indicated by average day lengths of more than 15 hours during shooting phase, which is necessary to trigger the effect of *Ppd-H1*^37^. In Dubai, where day length is shorter with less than 11 hours during shooting phase and more homogeneous across the year, no significant *Ppd-H1* effect on plant developmental traits was detected. In Al-Karak, we observed strong *Ppd-H1* effects, although this location is more than 2,000 km closer to the equator than Halle. Although the difference in day length between Al-Karak and Dubai seems negligible, a significant *Ppd-H1* effect on plant developmental traits was only present in Al-Karak. Therefore, we assume that an unknown interaction between day length and additional environmental cues, for instance temperature or precipitation, is necessary to trigger environment-dependent plant development effects of *Ppd-H1*^34,35,64^.

**Figure 3.**
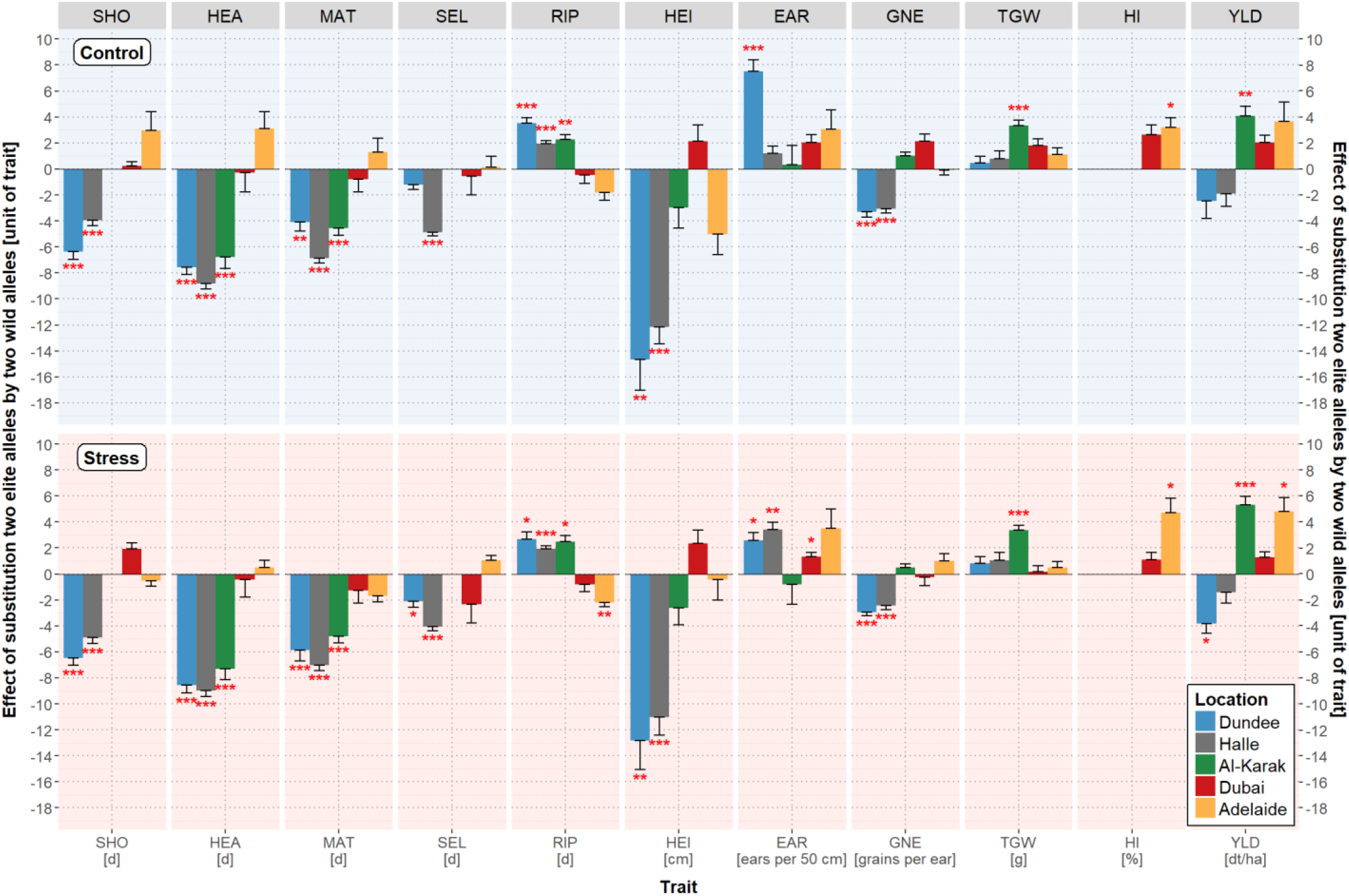
Estimates of *Ppd-H1* wild allele effects on plant developmental and yield-related traits. The trait names are given in the grey rectangles above each subplot and at the bottom where, in addition, the units of the traits are indicated. Trait abbreviations are listed in Supplementary Table S3. The color of the bars represents the location, blue for Dundee, grey for Halle, green for Al-Karak, red for Dubai and yellow for Adelaide. *Ppd-H1* wild allele effects under control and stress treatments are depicted with a bright blue (top) and a bright red background (bottom), respectively. Statistically significant wild allele effects are indicated by red asterisks above or below the bars with *P* < 0.05 = *, *P* < 0.01 = ** or *P* < 0.001 = ***. The height of the bars indicates the size of the *Ppd-H1* wild allele effect, obtained by calculating the difference between the mean performances of HEB-YIELD lines carrying two wild alleles versus two elite alleles.

In HEB-YIELD, *Ppd-H1* acted in a location-specific manner on yield-related traits. The most pronounced *Ppd-H1* effect was present in Al-Karak where the day length-sensitive wild barley allele was associated with an increase of grain yield by 4.1 dt/ha (+15 %) and 5.3 dt/ha (+30 %) under control and drought stress conditions, respectively (Figure 3; Supplementary Table S8a). The yield effect may be explained through pleiotropic effects of the wild barley *Ppd-H1* allele, which shortened the overall growing season, increased the period of grain filling (RIP) and increased grain size (TGW). A tendency of the *Ppd-H1* wild barley allele towards enhanced grain yields was also observed in Dubai and Adelaide, however, only significant in Adelaide under drought stress (+4.8 dt/ha = +29 %). Usually, the location-specific effects of *Ppd-H1* on yield-related traits are in agreement with the preferred length of the growing period. At those locations where earliness is beneficial, the responsive wild allele of *Ppd-H1* exerted increasing effects on yield-related traits, for example in Al-Karak, where early plants escaped higher temperatures and terminal drought at the end of the growing season. On the other hand, where lateness is preferable to achieve higher yields, the elite barley *ppd-H1* allele increased yield-related traits, for example, in Dundee and Halle. At these locations late HEB-YIELD lines benefited from the extended growing period since the environmental conditions supported plant growth under suitable conditions.

#### Sdw1

*Sdw1* belongs to the group of so-called semi-dwarfing genes^40^, which are responsible for yield elevations during the ‘Green Revolution’^65^. Wild barley accessions possess the functional and dominant *Sdw1* allele, a gibberellic acid 20 oxidase (*GA20ox*) gene, which promotes plant growth. In contrast, the recessive, GA-deficient *sdw1* allele^66,67^ is present in barley cultivars like Barke, causing a semi-dwarf phenotype. Several studies have shown that semi-dwarfs exhibit reduced plant height, late maturity, increased tiller numbers and an improved harvest index, ultimately resulting in elevated grain yields^40,68,69^.

The reported pleiotropic effects of *Sdw1* are also supported by HEB-YIELD field data (Figure 4; Supplementary Table S8a). Throughout plant development, we detected an accelerating effect of the wild barley *Sdw1* allele in HEB-YIELD, accelerating grain maturity by 4.0 to 8.9 days in Dundee, Al-Karak, Dubai and Adelaide, compared to the semi-dwarfing allele of Barke. Most striking was the pronounced delay of development in Adelaide under the control condition (precipitation=484 mm), with up to 13 days for SHO. Whereas under stress (precipitation=159 mm) the effects were on a similar level as in the other locations. The Adelaide effect might be explained by different environmental cues between the two years, resulting from the earlier sowing date and the prolonged growing period of 50 days in 2016. So far, there is no evidence that day length or precipitation affects the function of *Sdw1*^40,67^. However, a wheat survey under controlled conditions already reported that temperature can modify GA dependent responses, where elevated temperatures increase the abundance of GA^70^.

**Figure 4.**
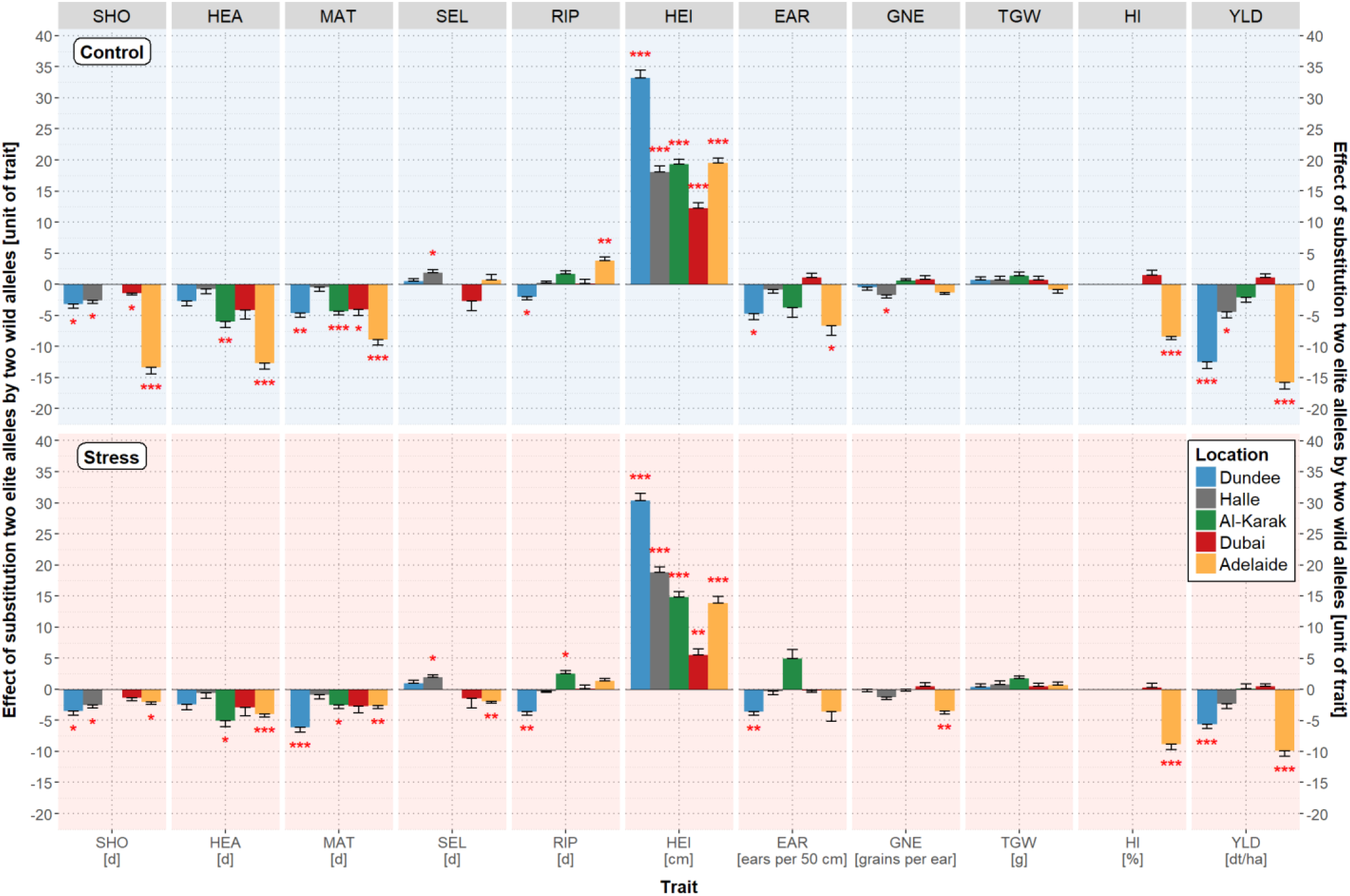
Estimates of *Sdw1* wild allele effects on plant developmental and yield-related traits. The trait names are given in the grey rectangles above each subplot and at the bottom where, in addition, the units of the traits are indicated. Trait abbreviations are listed in Supplementary Table S3. The color of the bars represents the location, blue for Dundee, grey for Halle, green for Al-Karak, red for Dubai and yellow for Adelaide. *Sdw1* wild allele effects under control and stress treatments are depicted with a bright blue (top) and a bright red background (bottom), respectively. Statistically significant wild allele effects are indicated by red asterisks above or below the bars with *P* < 0.05 = *, *P* < 0.01 = ** or *P* < 0.001 = ***. The height of the bars indicates the size of the *Sdw1* wild allele effect, obtained by calculating the difference between the mean performances of HEB-YIELD lines carrying two wild alleles versus two elite alleles.

The most prominent effect of semi-dwarfing genes is their control of plant architecture, in particular, plant height^40,71^. We confirmed this effect in HEB-YIELD since the wild barley allele increased plant height at all locations and under both treatments with a maximum increase of 33.2 cm in Dundee under control condition. The dominance of semi-dwarf genes in modern crop cultivars indicates their global importance for agriculture^65,72^.

This notion is also confirmed in HEB-YIELD where the Barke semi-dwarf allele was associated with an increase in grain yield (Figure 4; Supplementary Table S8a). In turn, the wild barley allele significantly reduced grain yield, for instance under control conditions in Dundee, Halle and Adelaide by up to 15.8 dt/ha. Under drought stress conditions in Adelaide, the Barke semi-dwarf allele revealed the strongest impact, accounting for 60 % of the final yield level. The observed yield increase may be attributed to an accumulation of several positive effects, including an extended growing period, more tillers, a higher harvest index and less lodging and head loss (Figure 3; Supplementary Table S8a).

#### Vrn-H1

In addition to the aforementioned photoperiod and GA dependent pathways, flowering time is also regulated through the vernalization pathway, where exposure to cold temperatures accelerates flowering^33,64,73^. In barley, the response to cold temperatures is mainly controlled by interaction of the two vernalization genes, *Vrn-H1*^74^ and *Vrn-H2*^75^. *Vrn-H2* acts as a strong repressor of flowering under long day conditions, preventing winter barley cultivars and wild barley accessions to flower during winter^75^. The expression of the *APETALA1 MADS-box gene Vrn-H1* is only induced after extended periods of cold exposure^76^, resulting in down-regulation of *Vrn-H2* and induction of flower initiation through direct binding of the Vrn-H1 protein to the promoters of *Vrn-H2* (repression) and *Vrn-H3* (activation)^77^. In spring barley cultivars like Barke the dominant *Vrn-H1* allele promotes flowering whereas the recessive winter barley and wild barley alleles delay flowering if cold exposure is imperfect.

In HEB-YIELD, *Vrn-H1* exhibited considerable effects on nearly every trait in Dubai and, to a lesser extent, in Adelaide (Supplementary Figure S3; Supplementary Table S8a). HEB-YIELD lines carrying the wild barley allele at this locus delayed flowering time and maturity by more than 10 days in Dubai. In Adelaide, pronounced effects on plant development were restricted to the control condition (i.e. the Adelaide growing period 2016). Most likely, this effect is caused by warmer temperatures and therefore less vernalization stimuli at the beginning of the growing season (Supplementary Figure S1 and S2; Supplementary Table S9). However, the late development effect of the wild barley *Vrn-H1* allele in Adelaide diminished during cultivation from +12 days at shooting, +6 days at flowering to, finally, +4 days at maturity. This tendency was also present in Halle and Al-Karak, although on a much lower level. In contrast, the late development effect of the wild barley *Vrn-H1* allele remained stable throughout plant cultivation in Dubai. This may be because the temperature in Dubai never reached a vernalization-triggering level. In Dubai, the HEB-YIELD lines possessing a wild barley winter allele at *Vrn-H1* thus responded to the lack of vernalization with a late plant development.

In addition to its developmental effects the wild barley allele of *Vrn-H1* exerted significant reducing effects on all yield components of around 25 % in Dubai. Consequently, the final grain yield in Dubai was reduced by 3.2 dt/ha under control conditions, which corresponds to 37 % of the total yield. At locations where earliness is the preferred breeding goal and vernalizing conditions are rare, the use of the dominant elite barley allele of *Vrn-H1* is highly recommended.

#### Vrn-H3

As mentioned before, the expression level of *Vrn-H1* increases with exposure to cold temperatures, resulting in flower induction through repression of *Vrn-H2* and activation of *Vrn-H3*^29,77^. *Vrn-H3* corresponds to the *HvFT1* gene, which is an orthologue of the Arabidopsis *FT* gene, the so called ‘florigen’^33,78,79^. *Vrn-H3* has a central role in flower induction integrating photoperiod and vernalization signals^79^. In barley, mutations in the first intron of the *VRN-H3* gene differentiate plants with respect to spring and winter growth habit. The dominant spring allele induces early flowering whereas the recessive winter barley allele delays flowering^80^.

HEB-YIELD field data validated the role of *Vrn-H3* on plant development throughout the whole growing period in all locations except from Dubai (Supplementary Figure S4; Supplementary Table S8a). The wild barley allele of *Vrn-H3* slowed down plant development between 2.2 and 6.6 days. Generally, winter genotypes are characterized by carrying a recessive *Vrn-H3* allele, which displays a reduced expression^80^. Most wild barleys possess a winter type^62^ and probably harbor a recessive v*rn-H3* allele, which explains the decelerating developmental effects.

Although *Vrn-H3* plays an important role for plant development, we identified only weak, mostly non-significant, impacts on yield-related traits. Only in Al-Karak, the wild allele showed significant reducing effects on grain number per ears (under both treatments) and on grain yield under drought stress (−3.3 dt/ha).

### The best wild barley HEB-YIELD lines match the yield performance of high-yielding local check cultivars

The usefulness of wild accessions, related to crop species has been proposed and demonstrated frequently^14,15,81^. Wild barley accessions, in particular *H. v.* ssp. *spontaneum,* the progenitor of cultivated barley have been used to improve disease resistance^24,82^ and abiotic stress tolerance^22,69,83,84^, as well as plant developmental traits^18,20,25^ and quality traits^82,85,86^. The successful use of wild relatives to increase grain yield of barley has not been reported frequently, some exceptions are available^69,87,88^. This may be because of the negative impacts of linked deleterious wild alleles, a phenomenon generally referred to as ‘linkage drag’^89^. The HEB-YIELD lines offer the possibility to estimate potentially positive wild allele effects in an adapted genetic background, since they are embedded through backcrossing into the modern elite barley cultivar Barke. In addition, the elite genetic background enables the direct use of HEB-YIELD lines in barley breeding programs.

Based on our two-year field trials we identified five high yielding HEB-YIELD lines, which showed high grain yield performance, comparable to the recurrent elite parent Barke, across the tested locations. These HEB-YIELD lines are 01_132, 01_104, 10_184, 10_173 and 05_043 (Supplementary Figure S5; Supplementary Tables S10a-S10c). They possessed higher grain yields than Barke in Al-Karak (except HEB_10_184). In addition, HEB-YIELD lines 01_132 and 10_184 surpassed the Barke grain yield in Dundee under both stress and control treatments. Furthermore, we identified HEB-YIELD lines, which reached or surpassed the yield level of locally adapted check cultivars (Supplementary Figure S5; Supplementary Tables S10a-S10c). These HEB lines are 10_184 and 01_132 in Dundee (Supplementary Figure S6), 01_132 and 01_104 in Halle (Supplementary Figure S7), 05_043 and 10_173 in Al-Karak (Figure 5), 15_082 and 06_116 in Dubai (Supplementary Figure S8) and 10_184 & 01_132 in Adelaide (Supplementary Figure S9). For instance, HEB_01_132 surpassed the grain yield of the established local check cultivar ‘Navigator’ under stress treatment in Adelaide. In addition, under both treatments it was comparable to ‘Compass’ and ‘La Trobe’, which have become the dominant commercial cultivars in South Australia. HEB_01_132 also surpassed the grain yield of the local check ‘58/1 A’ under control treatment in Dubai, indicating that this line may be directly suited for cultivation in the respective environments. Likewise, HEB_05_043 and HEB_10_173 outperformed the check cultivar ‘Rum’ in Al-Karak under drought stress.

**Figure 5.**
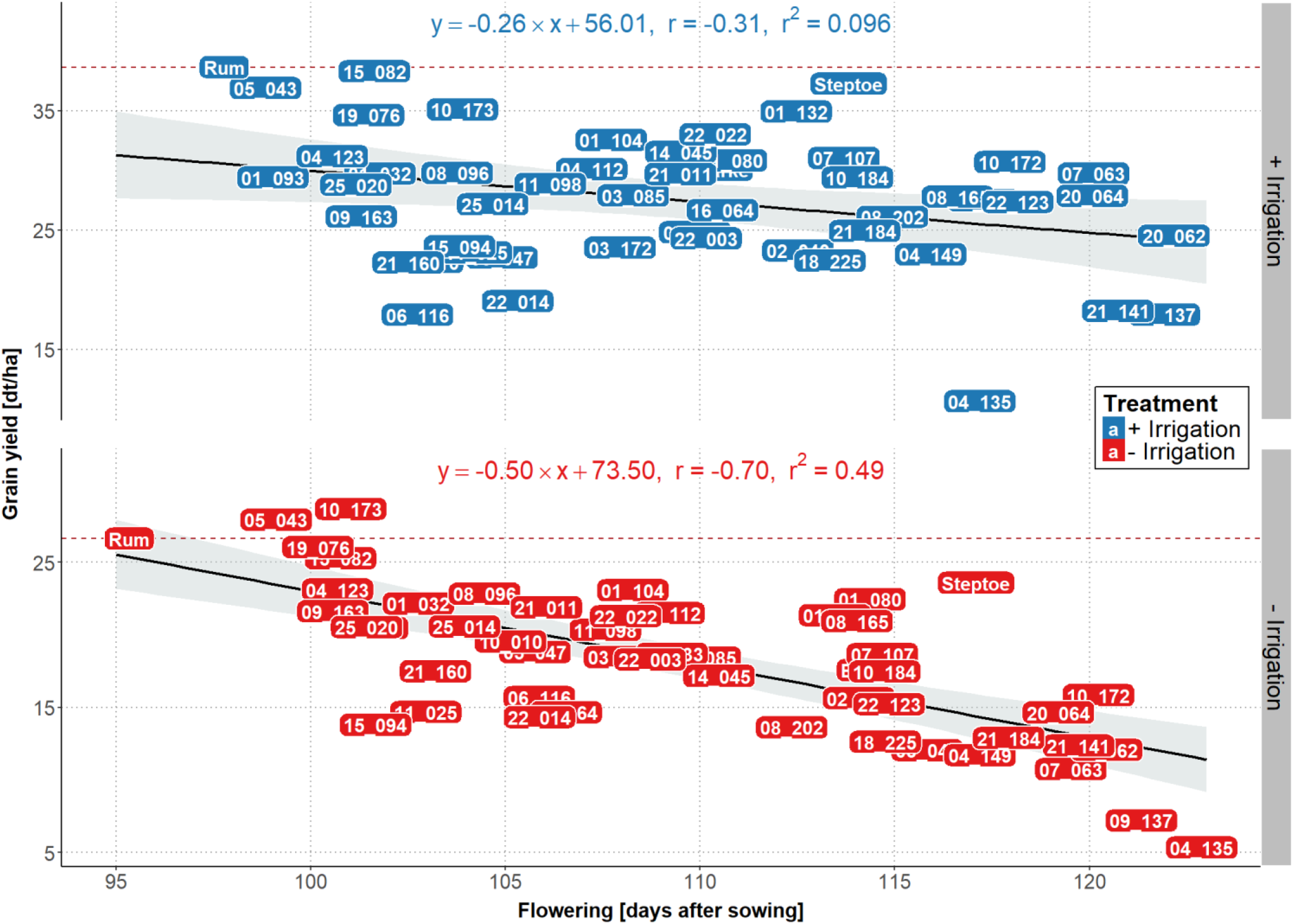
Regression of grain yield on flowering in Al-Karak. The yield levels of the 48 HEB-YIELD lines plus checks are depicted as a function of flowering time, separately for control (blue labels) and stress (red labels) treatments. The yield level of the local check cultivar ‘Rum’ is indicated by a dashed red line. On top of each subplot the linear regression equation, the Pearson’s correlation coefficient (*r*) and the coefficient of determination (*r*^*2*^) are indicated.

Interestingly, HEB-YIELD lines adapted different yield formation strategies at each test location. Compared to the local check cultivars, HEB-YIELD lines had increased numbers of ears (EAR) at Al-Karak, increased thousand grain weights at Dubai in almost all cases, and increased grain numbers per ear under stress at Dundee and Adelaide in many cases (Supplementary Figures S10-S12). This offers the possibility of achieving future yield improvements following a location-specific adaptation route.

The challenges of climate change demand that cultivars need to re-adapt to changing environmental conditions, for instance shorter growing seasons, higher average temperatures during cultivation and more frequently occurring drought periods^26,50,90^. HEB-YIELD lines exhibited a high phenological variation. For instance, flowering time exhibited a range of 49 days in Dubai and 36 days in Adelaide (Supplementary Table S5), which offers the potential to use this variation to adapt new cultivars to changing environmental conditions by backcrossing favorable HEB-YIELD donor lines with locally adapted elite cultivars. In those areas where drought and heat affect plant development and grain maturation, early maturing lines like HEB_05_043, HEB_15_082 or HEB_10_173 may be beneficial because of their fast development (Figure 5; Supplementary Figure S9; Supplementary Table S10a). Moreover, the increased tiller capacity of HEB-YIELD lines may be promising to achieve an improved canopy cover, reducing moisture losses^26,49^ and to increase biomass yield. The latter trait may be high value in the eastern part of the Mediterranean basin where straw and grains of barley are mainly used for animal feeding^91,92^.

In the past, Australian varieties followed a strong focus on earliness but changes in agricultural practices have resulted in earlier sowing dates and thus an extended growing season. The earlier sowing allowed later genotypes to benefit from a longer growing period, enabling HEB-YIELD lines HEB_03_085, HEB_10_184, 20_064 and HEB_04_135 to surpass the grain yield of the local check cultivar ‘Navigator’ in Adelaide under control condition (Supplementary Figure S9; Supplementary Table S10a). We also identified HEB-YIELD lines that performed quite well in the high yielding environments of Dundee and Halle. Here, lines HEB_01_104, HEB_01_132 and HEB_10_184 accomplished reasonable yields. HEB_10_184, for instance, achieved a maximum grain yield of 74.0 dt/ha under control condition in Dundee, which was almost on par with the local check cultivar ‘Odyssey’ (−0.3 %) and 5.1 % higher than the recipient cultivar Barke. This finding indicates that in future wild barley germplasm may be used to improve grain yield, for instance, by extend the growing period, as temperature and precipitation in Scotland are adequate to support late genotypes (Supplementary Figures S1 and S2; Supplementary Tables S9a and S10a).

## Conclusion

It is expected that the impact of climate change necessitates the adaptation of our established crop cultivation systems to harsher environmental conditions^26,90^. Stress avoidance is one promising approach to increase stress tolerance. We explored this relationship by studying the wild barley-derived model population HEB-YIELD in a field experiment, ranging from Dundee in Scotland to Adelaide in South Australia, where the effects of nitrogen deficiency, drought and salinity on plant development and yield-related traits were investigated.

Our findings confirm the crucial relationship between flowering time, plant development and grain yield^93^. The exact timing of the switch from vegetative to reproductive growth under favorable conditions^32^, the length of the growing period and the duration of the sub-phases of plant development are crucial to secure yield under abiotic stress conditions. We suggest that adjusting plant development may be a promising breeding strategy to cope with abiotic stresses. To optimize breeding programs, it is thus advisable to first predict the environment-dependent impact of flowering time genes on yield formation and then to select locally advantageous alleles for sustainable crop improvement.

Our HEB-YIELD data indicate that wild germplasm may serve as a resource to increase genetic diversity^14,20,22^ and to enable the above mentioned adaptation to abiotic stresses, through selection of early or late development alleles of known major flowering time genes, e.g. *Ppd-H1, Sdw1, Vrn-H1* and *Vrn-H3*. We showed that allelic variants of these flowering time genes strongly react to environmental cues. This information can be used to design novel breeding strategies such as precise backcrossing of suitable developmental genes into regionally adapted cultivars. Our data also provide evidence that wild barley germplasm may be useful to improve yield in low-yielding environments, for instance, in the Middle East, as well as in high-yielding environments, for instance, in Northern and Central Europe. This knowledge may be transferred to related crop species like wheat and rice to secure the rising global food demand for cereals.

## Materials and Methods

### Plant material

HEB-YIELD, a subset of the wild barley nested association mapping (NAM) population Halle Exotic Barley-25 (HEB-25^18^), was used in yield trials. HEB-25 originated from crossing 25 diverse wild barley accessions (*Hordeum vulgare* ssp. *spontaneum* and *H.v.* ssp. *agriocrithon*) with the German spring barley elite cultivar Barke (*Hordeum vulgare* ssp. *vulgare*). HEB-25 comprises 1,420 BC_1_S_3_ derived lines (backcrossed with Barke) grouped into 25 families (for more details see Maurer *et al*.^18^).

The HEB-YIELD subset consists of 48 HEB-25 lines that were selected from HEB-25 to ensure the absence of brittleness and a good threshability enabling accurate yield estimation in field trials. In addition, the final HEB-YIELD lines were selected to independently segregate at four major flowering time loci, which exhibited major plant developmental effects in HEB-25: *Ppd-H1, Sdw1, Vrn-H1* and *Vrn-H3*^18,20,25^.

### Genotypic data

The complete HEB-25 population was genotyped in generation BC_1_S_3_ using the barley Infinium iSelect 9k SNP chip (see^18^). The diagnostic markers i_BK_16, i_12_30924, i_11_10705 and i_12_10218, co-segregating with the four flowering time genes *Ppd-H1, Sdw1, Vrn-H1* and *Vrn-H3,* respectively, were used for selection of HEB-YIELD lines carrying homozygous elite versus homozygous wild barley alleles (Supplementary Table S1).

### Field trials

The HEB-YIELD population was grown at five locations worldwide during two years (2015 and 2016), resulting in ten environments. The locations are (from north to south): Dundee (United Kingdom; 56°28′53.71″N 3°6′35.17″W), Halle (Germany; 51°29′46.05″N 11°59′29.58″E), Al-Karak (Jordan; 31°16′34.03″N 35°44′24.94″E), Dubai (United Arab Emirates; 25°5′44.40″N 55°23′24.48″E) and Adelaide (Australia; 35°19′18.5″S 138°53′07.5″E). A detailed description for each location is given in Supplementary Table S2a. The full set of 48 HEB-YIELD lines was cultivated at each location except in Adelaide. Due to lack of seeds, in Adelaide only 34 and 47 HEB-YIELD lines were cultivated in 2015 and 2016, respectively (Supplementary Table S2d). At each location, additional local check cultivars were cultivated, for example: ‘Odyssey’ (Limagrain, 2011) in Dundee, ‘Quench’ (Syngenta, 2006) in Halle, ‘Rum’ (CIMMYT, 1986) in Al-Karak, ‘58/1 A’ (ICBA, 2002) in Dubai and ‘Navigator’ (University of Adelaide, 2012) in Adelaide.

At each location, a control treatment and a site-specific stress treatment was applied. Stress treatments were nitrogen deficiency in Dundee and Halle, drought stress in Al-Karak, salt stress in Dubai and drought stress in Adelaide (see Supplementary Table S2c). Therefore, lines of the stress treatment received no nitrogen fertilizer in Dundee and Halle, no drip irrigation in Al-Karak and a saline water drip irrigation in Dubai. In Adelaide, only one treatment was applied per season due to lack of seeds. In this case, the two contrasting seasons represented the treatments where 2015 was regarded as the drought stress treatment with only 159 mm precipitation during the growing period and 2016 as the control treatment with 484 mm precipitation.

On average, each HEB-YIELD line was replicated three to four times per treatment. A randomized complete block design was chosen as test design for the trials, with the exception of Dubai and Adelaide where a completely randomized design within each treatment was applied. The trials were conducted in adaptation to local practices regarding tillage, fertilization and pest management. Additional information on plant cultivation is provided in Supplementary Table S2b.

### Phenotypic data

Eleven developmental and yield related traits were investigated. A description of where and how each trait was measured is given in Supplementary Table S3.

### Statistical analyses

All statistical analyses were carried out with SAS 9.4 (SAS Institute Inc., Cary, NC, USA)^41^. Variance components (defined as random) were estimated with *PROC VARCOMP* and broad sense heritabilities (h^2^) for each trait within locations and treatments were calculated across years following the formula:

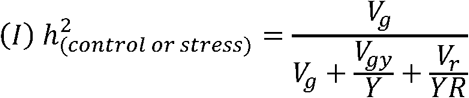

where

*V*_*g*_: = genotypic variance
*Y*: = number of years
*V*_*gy*_: = genotype by year interaction variance
*R*: = number of replications
*V*_*r*_: = error variance

For traits analyzed in a single year, repeatability (rep) was calculated following the formula:

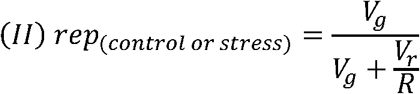

The analysis of variance (ANOVA) across locations was calculated with *PROC MIXED* to test for the presence of genotype, location and year effects. For this purpose, the main effects (genotype, location and year), as well as their corresponding interaction effects were treated as fixed effects in the following model:

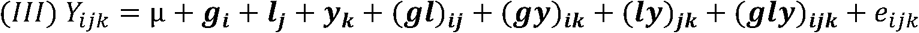

where

*Y*_*ijk*_: = observed phenotype of the *i*th genotype in the *j*th location and the *k*th year
µ: = Intercept
***g***_***i***_: = effect of the *i*th genotype
***l***_***j***_: = effect of the *j*th location
***y***_***k***_: = effect of the *k*th year
(***gl***)_***ij***_: = interaction effect between the *i*th genotype and the *j*th location
(***gy***)_***ik***_: = interaction effect between the *i*th genotype and the *k*th year
(***ly***)_***jk***_: = interaction effect between the *j*th location and the *k*th year
(***gly***)_***ijk***_: = interaction effect between the *i*th genotype, the *j*th location and the *k*th year
*e*_*ijk*_: = residual/error of *y*_*ijk*_

Fixed effects are written in bold

Best linear unbiased estimators (BLUEs) were estimated using the *PROC MIXED* procedure. The BLUEs for each HEB-YIELD line were computed across years and for each treatment level and location separately. Genotype and treatment were modelled as fixed effects and year as a random effect:

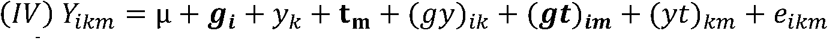

where

*Y*_*ikm*_: = observed phenotype of the *i*th genotype in the *k*th year and the *m*th treatment
µ: = Intercept
***g***_***i***_: = effect of the *i*th genotype
*y*_*k*_: = effect of the *k*th year
***t***_***m***_: = effect of the *m*th treatment
(*gy*)_*ik*_: = interaction effect between the *i*th genotype and the *k*th year
(***gt***)_***im***_: = interaction effect between the *i*th genotype and the *m*th treatment
(*yt*)_*km*_: = interaction effect between the *k*th year and the *m*th treatment
*e*_*ikm*_: = residual/error of *y*_*ikm*_

Fixed effects are written in bold

For location Adelaide BLUEs were calculated within years and the model was restricted to a fixed effect of genotype.

Pearson correlation coefficients (r) between trait BLUEs were calculated via *PROC CORR*. Furthermore, to test for significant treatment effects a simple t-test (*PROC TTEST*) and an ANOVA within locations were performed (*PROC MIXED*). The ANOVA model included the main effects (genotype, treatment and year) and their corresponding interaction effects as fixed effects (comparable to model III). In addition, a one-factorial ANOVA was computed to test for significant location effects within treatments where only the main effect (location) was included, followed by a Tukey test (*PROC GLM*).

Performance of the HEB-YIELD lines was compared to an adapted check cultivar from the corresponding location (see field trials above) by conducting a Dunnett test^42^ (*PROC MIXED*). To enable an easier comparison between the lines the relative performance (RP) was calculated as:

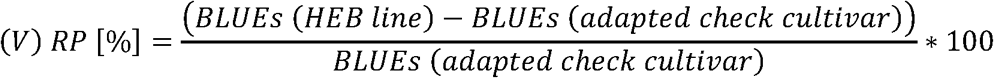

To check for significance and estimate effects of the four flowering candidate genes *Ppd-H1, Sdw1, Vrn-H1* and *Vrn-H3*, a simple linear regression model (*PROC GLM*) was fitted for each candidate gene applying BLUEs across years. Each model included a single locus-specific SNP mentioned above, modeled as a quantitative variable representing the wild allele dosage^20^.

All figures were created using R (3.4.2)^43^ with the package ggplot2 (2.2.1)^44^.

## Supporting information

## Acknowledgments

This work was financially supported by the German Research Foundation (DFG) via the priority program 1530: ‘Flowering time control - from natural variation to crop improvement’ with grants Pi339/7-1 and Pi339/7-2) and via ERA-NET for Coordinating Action in Plant Sciences (ERA-CAPS) grant Pi339/8-1. Funding from King Abdullah University of Science and Technology (KAUST) is also gratefully acknowledged. We are grateful to a multitude of research assistants, located at the James Hutton Institute in Dundee, the Martin Luther University in Halle, the International Center for Agricultural Research in the Dry Areas and NCARE in Al-Karak, the International Center for Biosaline Agriculture in Dubai and at the University of Adelaide for their excellent technical support in conducting the global field trials.

## Author Contribution Statement

MW conducted the field trials in 2015 and 2016 in Halle, gathered and analyzed the phenotypic data of all five test locations, created the figures, and drafted the manuscript. AM planned the field trials in 2015 and 2016 in Halle, supported the analysis of the phenotypic data and drafted the manuscript. AF, HB and WT planned and conducted the field trials in 2015 and 2016 in Dundee. MB and AA planned and conducted the field trials in 2014/15 and 2015/16 in Al-Karak. MT and MS planned and conducted the field trials in 2014/15 and 2015/16 in Dubai. JE, AP and TM planned and conducted the field trials in 2015 and 2016 in Adelaide. KP acquired the funding, coordinated the collaboration between the project partners and drafted the manuscript.

## Supplementary information

### Supplementary tables

**Supplementary Table S1** “Genotype segregation of flowering time genes in HEB-YIELD”

**Supplementary Table S2** “Description of the experimental field trials”

**Supplementary Table S3** “List of scored traits”

**Supplementary Table S4** “Raw data of field trials”

**Supplementary Table S5** “Descriptive statistics and heritabilities”

**Supplementary Table S6** “ANOVA”

**Supplementary Table S7** “Pearson’s correlation coefficients”

**Supplementary Table S8** “Flowering time gene effects”

**Supplementary Table S9** “Climate data 2015 & 2016”

**Supplementary Table S10** “Dunnett’s test”

### Supplementary figures

**Supplementary Figure S1** “Climate graph for growing period 2014/15 presenting day length, temperature and precipitation per location”

**Supplementary Figure S2** “Climate graph for growing period 2015/16 presenting day length, temperature and precipitation per location”

**Supplementary Figure S3** “Estimates of *Vrn-H1* wild allele effects on plant development and yield-related traits”

**Supplementary Figure S4** “Estimates of *Vrn-H3* wild allele effects on plant development and yield-related traits”

**Supplementary Figure S5** “Relative grain yield performance of 48 HEB YIELD lines compared to a local check cultivar under control and stress treatments”

**Supplementary Figure S6** “Regression of grain yield on flowering in Dundee”

**Supplementary Figure S7** “Regression of grain yield on flowering in Halle”

**Supplementary Figure S8** “Regression of grain yield on flowering in Dubai”

**Supplementary Figure S9** “Regression of grain yield on flowering in Adelaide”

**Supplementary Figure S10** “Relative ear number performance of 48 HEB YIELD lines compared to a local check cultivar under control and stress treatments”

**Supplementary Figure S11** “Relative grain number per ear performance of 48 HEB YIELD lines compared to a local check cultivar under control and stress treatments”

**Supplementary Figure S12** “Relative thousand grain weight performance of 48 HEB YIELD lines compared to a local check cultivar under control and stress treatments”

