## Supplementary material for "Barley yield formation under abiotic stress depends on the interplay between flowering time genes and environmental cues"

### Supplementary figures – S1-S12

#### Outline

### Supplementary Figure S1. Climate graph for growing period 2014/15 presenting day length, temperature and precipitation per location

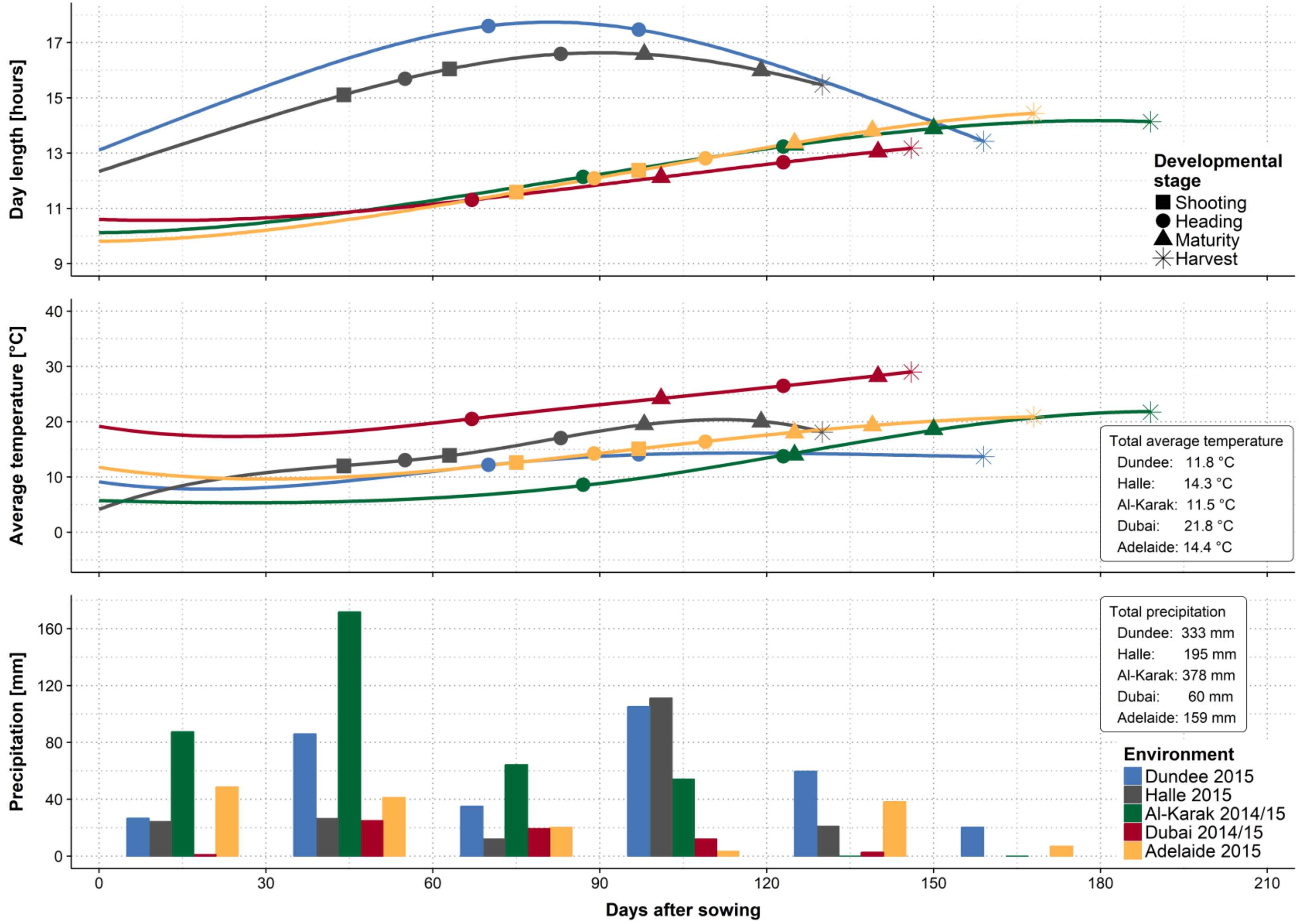

**Figure S1.** The upper plot illustrates day length in hours, the center plot the average temperature (i.e. minimum plus maximum temperature divided by two) in degree Celsius and the bottom plot the precipitation in mm per 30 days. The x-axis indicates days after sowing for all plots. The locations Dundee (DUN), Halle (HAL), Al-Karak (ALK), Dubai (DUB) and Adelaide (ADE) are illustrated in blue, grey, green, red and yellow, respectively. The developmental stages shooting, heading, maturity and harvest are indicated by square, circle, rectangle and star symbols, respectively. The first and the second appearance of the symbols specifies the first and the last occurrence of the stage in HEB-YIELD lines. The average temperature and precipitation during the growing period is displayed at the right border of the respective plot. Weather data were recorded at each field site.

#### Supplementary Figure S2. Climate graph for growing period 2015/16 presenting day length, temperature and precipitation per location

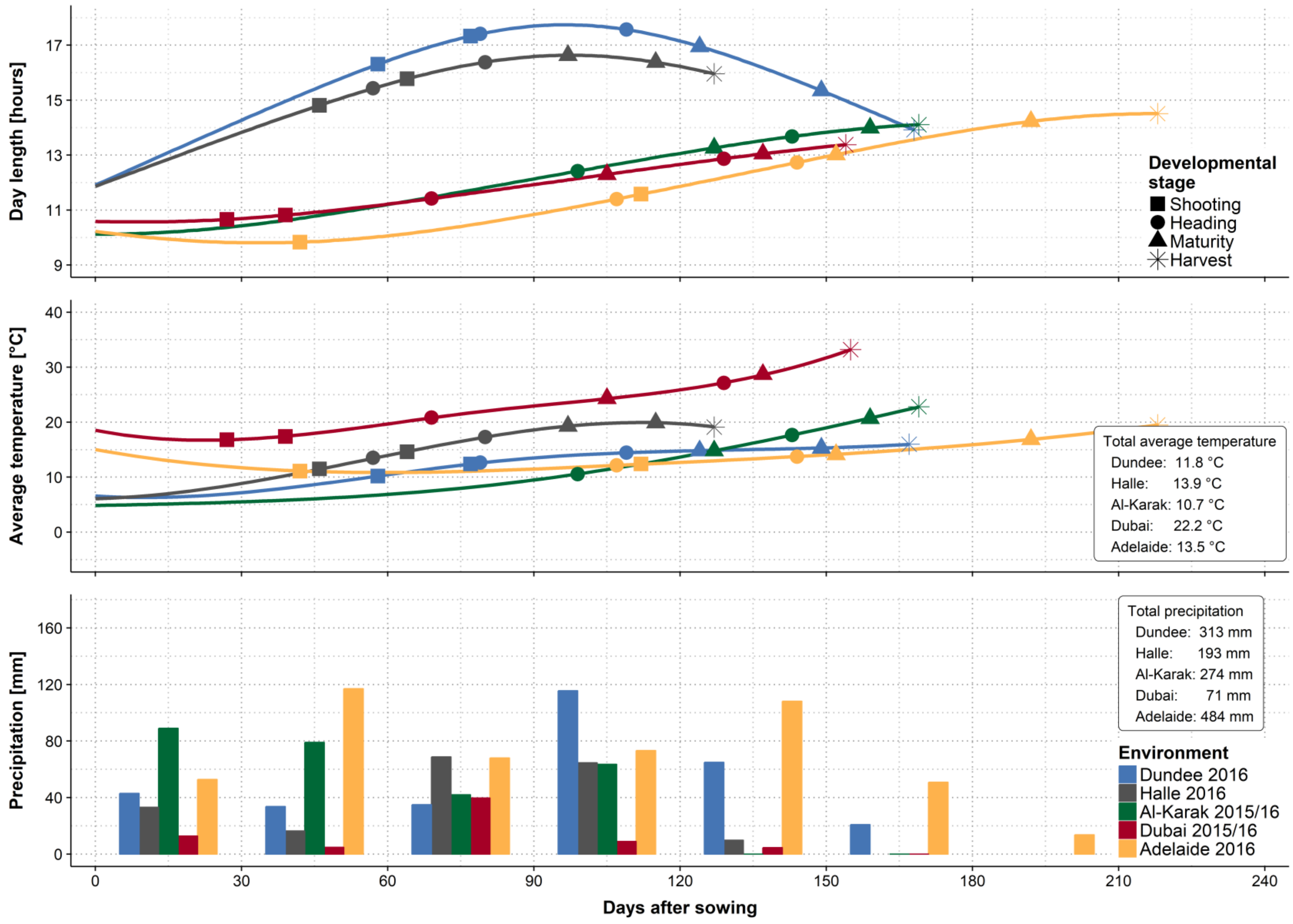

**Figure S2.** The upper plot illustrates day length in hours, the center plot the average temperature (i.e. minimum plus maximum temperature divided by two) in degree Celsius and the bottom plot the precipitation in mm per 30 days. The x-axis indicates days after sowing for all plots. The locations Dundee (DUN), Halle (HAL), Al-Karak (ALK), Dubai (DUB) and Adelaide (ADE) are illustrated in blue, grey, green, red and yellow, respectively. The developmental stages shooting, heading, maturity and harvest are indicated by square, circle, rectangle and star symbols, respectively. The first and the second appearance of the symbols specifies the first and the last occurrence of the stage in HEB-YIELD lines. The average temperature and precipitation during the growing period is displayed at the right border of the respective plot. Weather data were recorded at each field site.

### Supplementary Figure S3. Estimates of *Vrn-H1* wild allele effects on plant development and yield-related traits

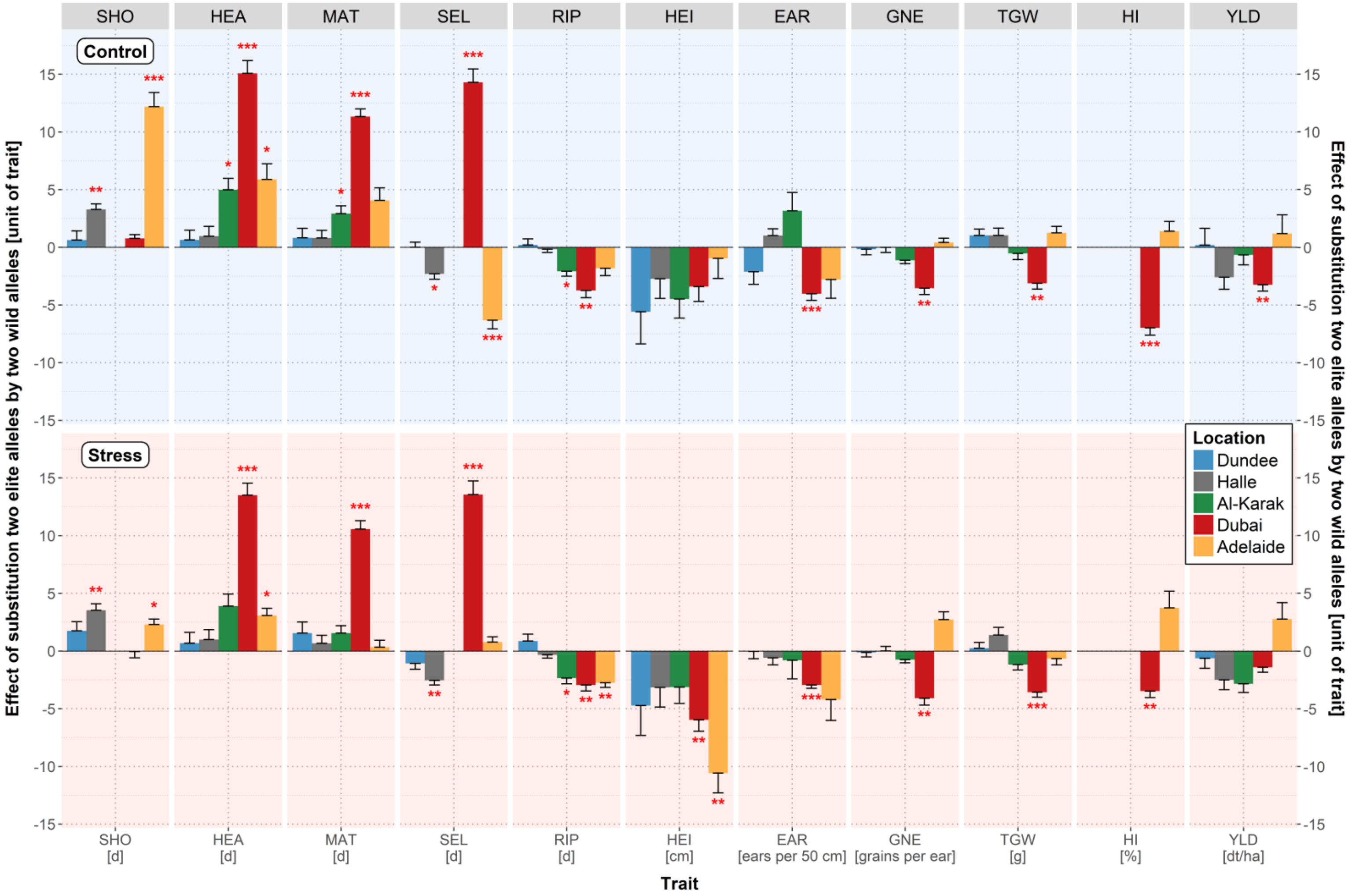

**Figure S3.** The trait names are given in the grey rectangles above each subplot and at the bottom where, in addition, the units of the traits are indicated. Trait abbreviations are listed in Supplementary Table S3. The color of the bars represents the location, blue for Dundee, grey for Halle, green for Al-Karak, red for Dubai and yellow for Adelaide. *Vrn-H1* wild allele effects under control and stress treatments are depicted with a bright blue (top) and a bright red background (bottom), respectively. Statistically significant wild allele effects are indicated by red asterisks above or below the bars with $P < 0.05$ = *, $P < 0.01$ = ** or $P < 0.001$ = ***. The height of the bars indicates the size of the *Vrn-H1* wild allele effect, obtained by calculating the difference between the mean performance of HEB-YIELD lines carrying two wild alleles versus two elite alleles.

### Supplementary Figure S4. Estimates of *Vrn-H3* wild allele effects on plant development and yield-related traits

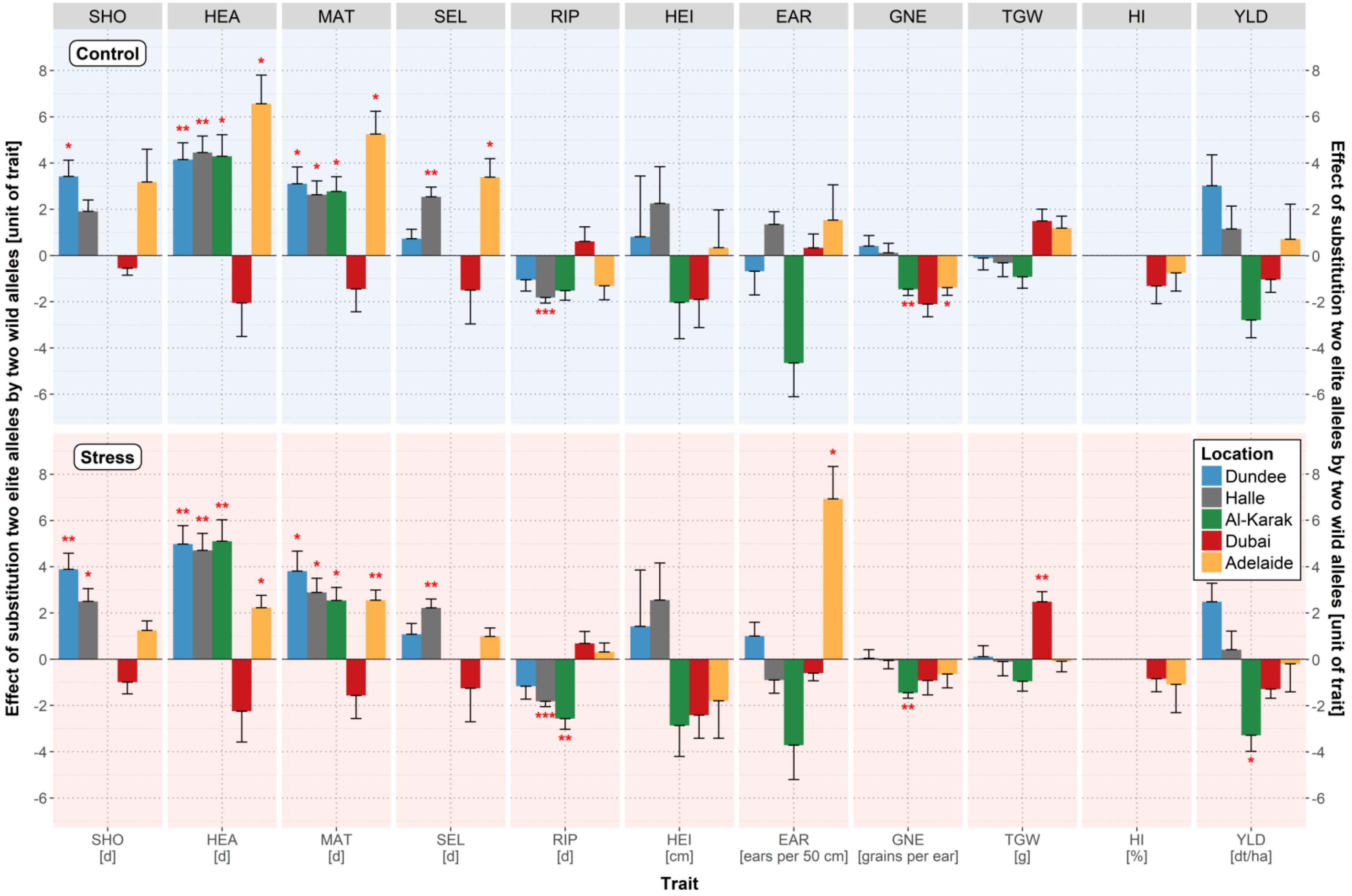

**Figure S4.** The trait names are given in the grey rectangles above each subplot and at the bottom where, in addition, the units of the traits are indicated. Trait abbreviations are listed in Supplementary Table S3. The color of the bars represents the location, blue for Dundee, grey for Halle, green for Al-Karak, red for Dubai and yellow for Adelaide. *Vrn-H3* wild allele effects under control and stress treatments are depicted with a bright blue (top) and a bright red background (bottom), respectively. Statistically significant wild allele effects are indicated by red asterisks above or below the bars with $P < 0.05$ = *, $P < 0.01$ = ** or $P < 0.001$ = ***. The height of the bars indicates the size of the *Vrn-H3* wild allele effect, obtained by calculating the difference between the mean performance of HEB-YIELD lines carrying two wild alleles versus two elite alleles.

### Supplementary Figure S5. Relative grain yield performance of 48 HEB YIELD lines compared to a local check cultivar under control and stress treatments

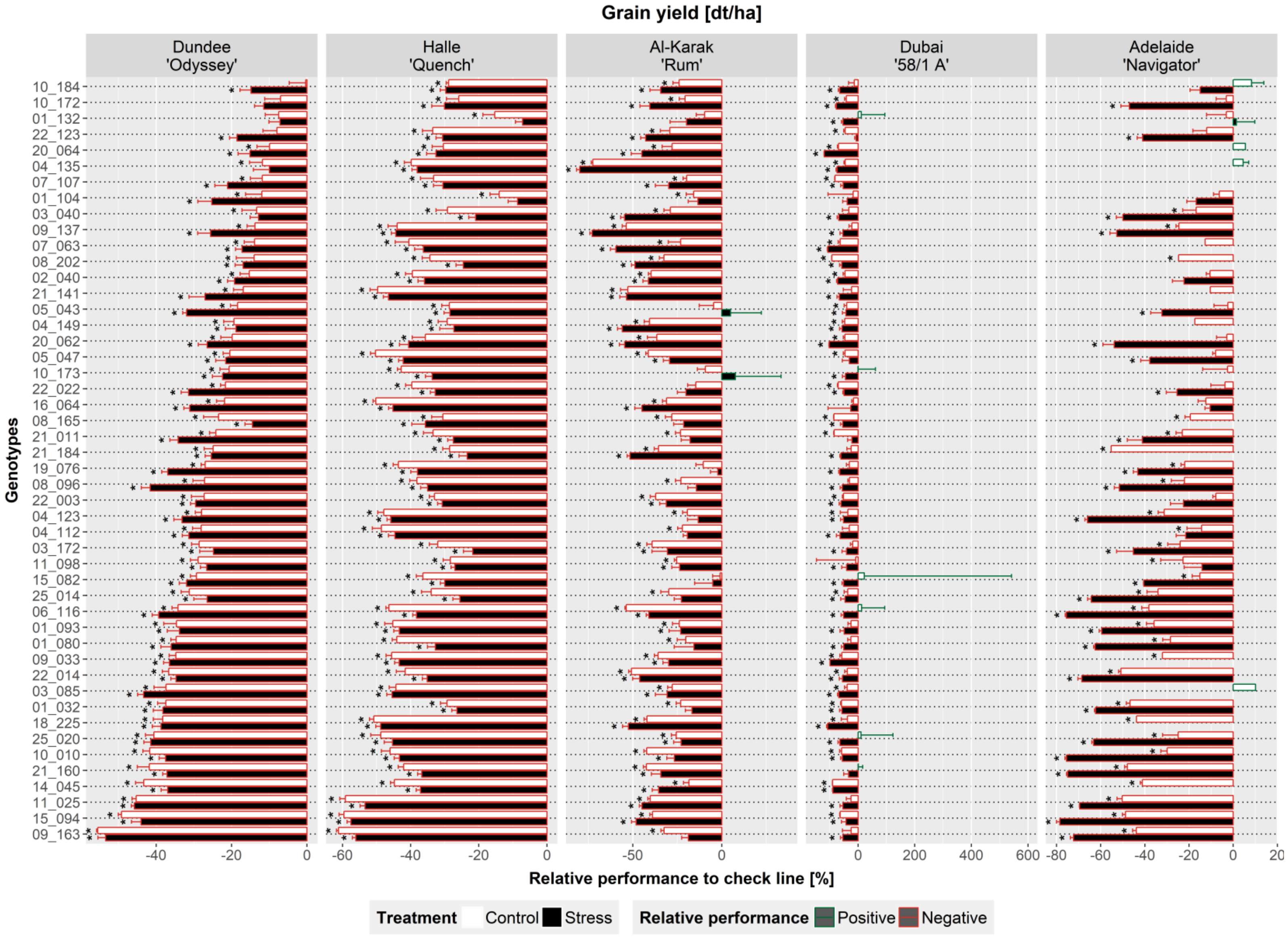

**Figure S5.** Location names and the respective check cultivar are indicated in the grey rectangles above each plot. On the x-axis, the relative performance (in %) of HEB-YIELD lines compared to the check cultivar is depicted. On the y-axis, the 48 HEB-YIELD lines are listed, sorted based on performance under control treatment in Dundee. The increase and decrease in performance relative to the check cultivar is indicated in red and green bars, respectively. Empty and filled bars indicate relative performance under control and stress treatments, respectively. Based on a Dunnett test, HEB-YIELD lines significantly ($P < 0.05$) deviating from the check cultivar are marked with black asterisks above the bars.

### Supplementary Figure S6. Regression of grain yield on flowering in Dundee

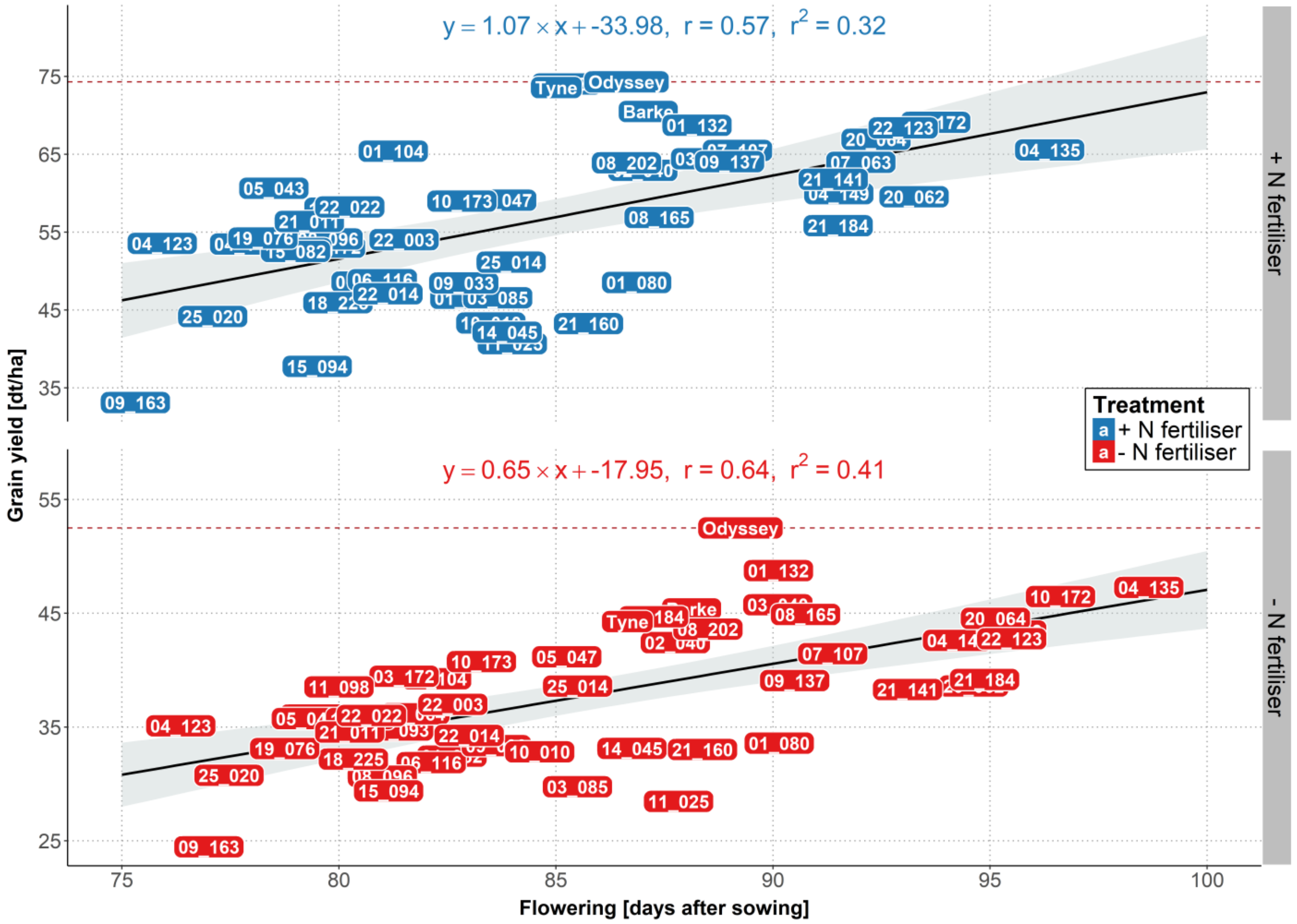

**Figure S6.** The yield levels of 48 HEB-YIELD lines plus checks are depicted as a function of flowering time, separately for control (blue labels) and stress (red labels) treatments. The yield level of the local check cultivar ‘Odyssey’ is indicated by a dashed red line. On top of each subplot the linear regression equation, the Pearson’s correlation coefficient (r) and the coefficient of determination ($r^2$) are indicated.

### Supplementary Figure S7. Regression of grain yield on flowering in Halle

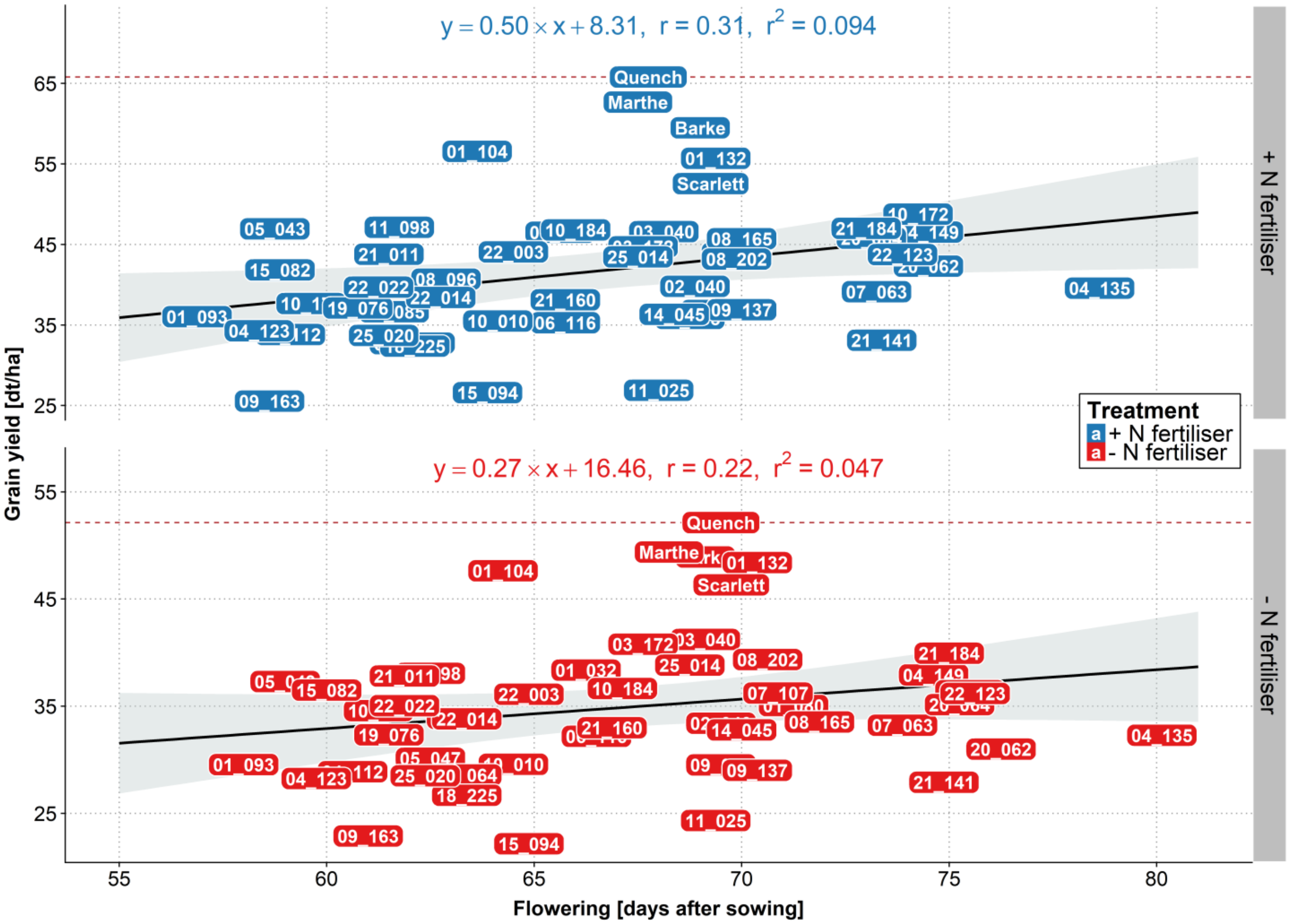

**Figure S7.** The yield levels of 48 HEB-YIELD lines plus checks are depicted as a function of flowering time, separately for control (blue labels) and stress (red labels) treatments. The yield level of the local check cultivar 'Quench' is indicated by a dashed red line. On top of each subplot the linear regression equation, the Pearson's correlation coefficient (r) and the coefficient of determination ($r^2$) are indicated.

### Supplementary Figure S8. Regression of grain yield on flowering in Dubai

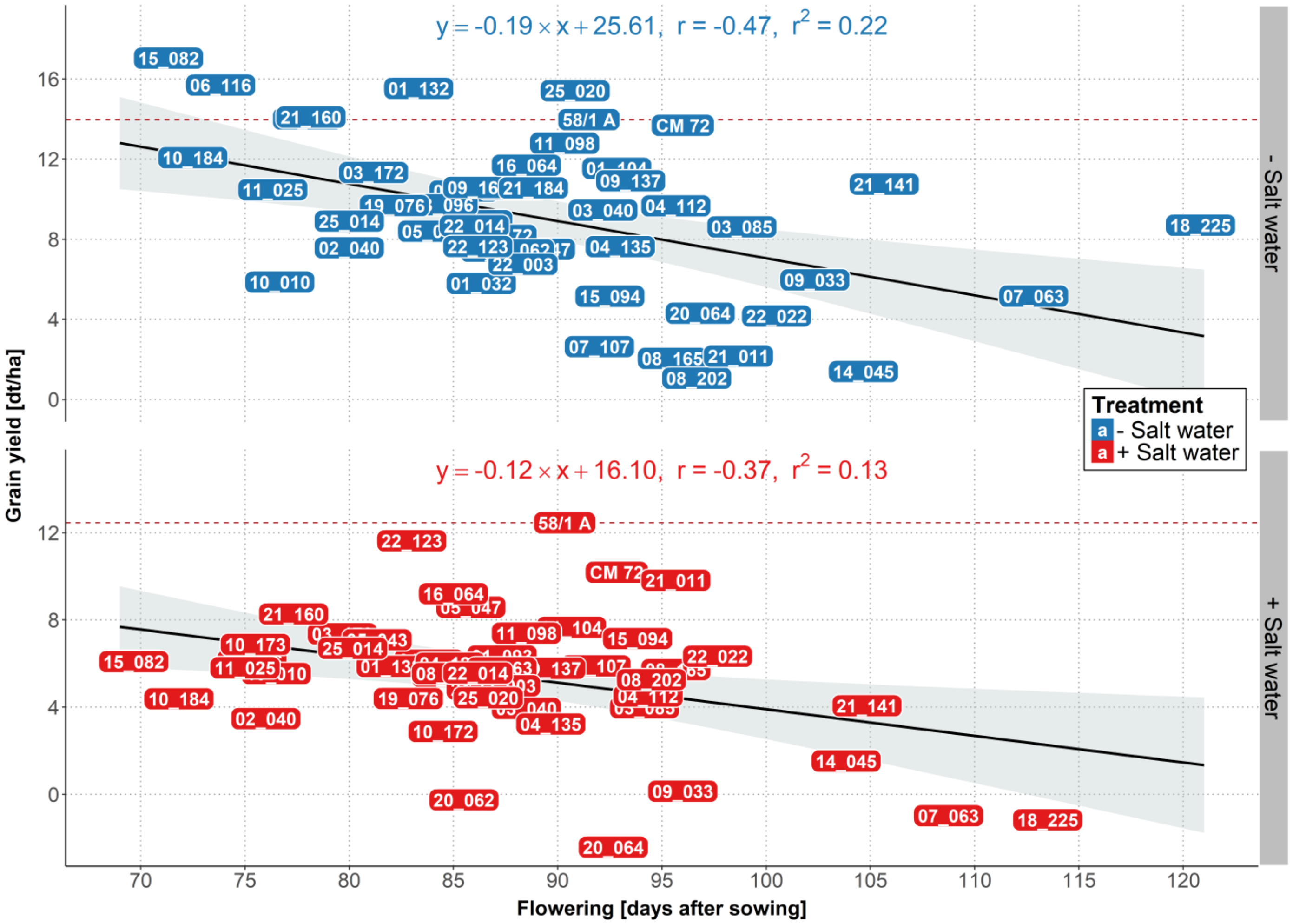

**Figure S8.** The yield levels of 48 HEB-YIELD lines plus checks are depicted as a function of flowering time, separately for control (blue labels) and stress (red labels) treatments. The yield level of the local check cultivar '58 1/A' is indicated by a dashed red line. On top of each subplot the linear regression equation, the Pearson's correlation coefficient (r) and the coefficient of determination ($r^2$) are indicated.

### Supplementary Figure S9. Regression of grain yield on flowering in Adelaide

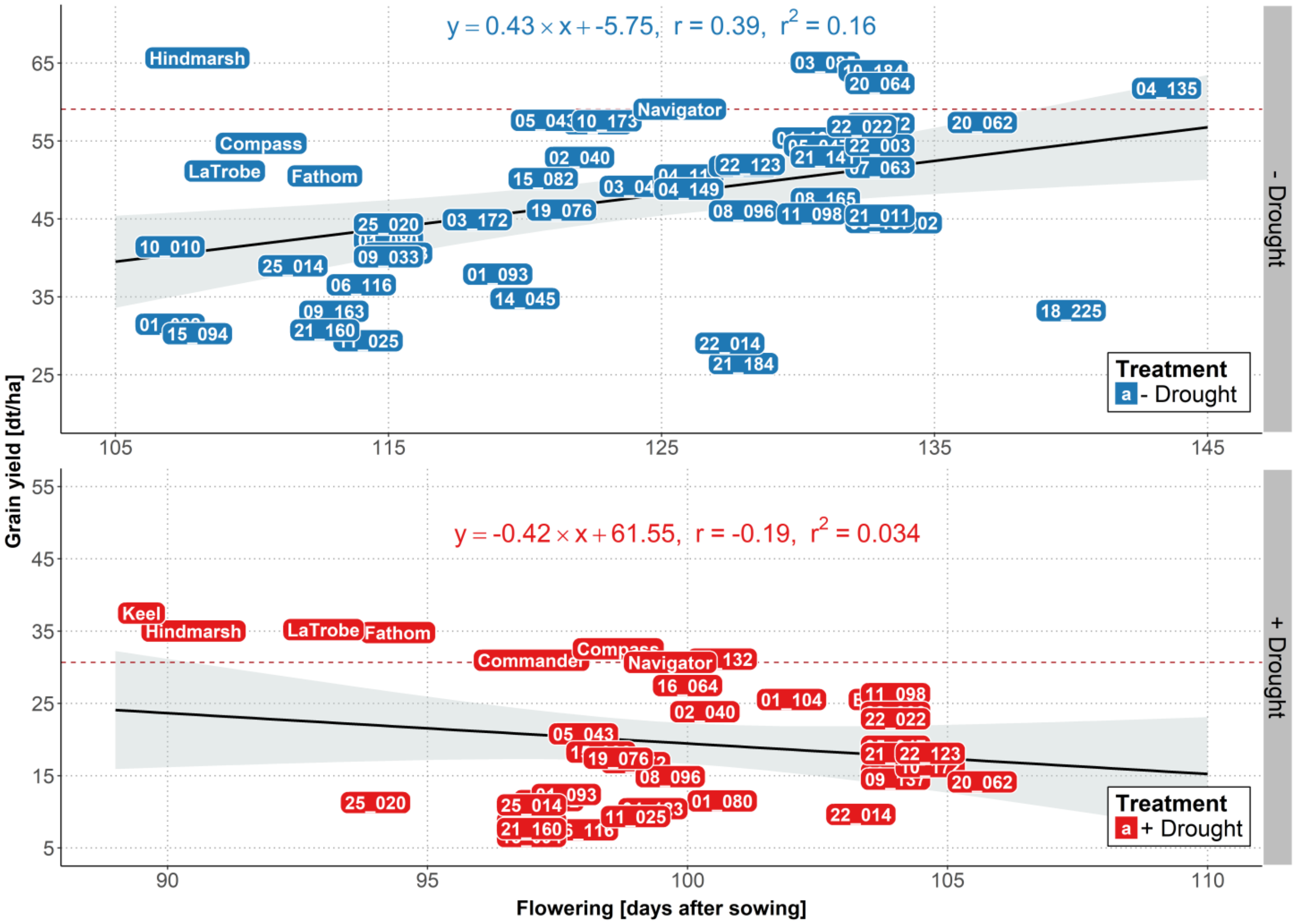

**Figure S9.** The yield levels of the 48 HEB-YIELD lines plus checks are depicted as a function of flowering time, separately for control (blue labels) and stress (red labels) treatments. The yield level of the local check cultivar 'Navigator' is indicated by a dashed red line. On top of each subplot the linear regression equation, the Pearson's correlation coefficient ($r$) and the coefficient of determination ($r^2$) are indicated.

#### Supplementary Figure S10. Relative ear number performance of 48 HEB YIELD lines compared to a local check cultivar under control and stress treatments

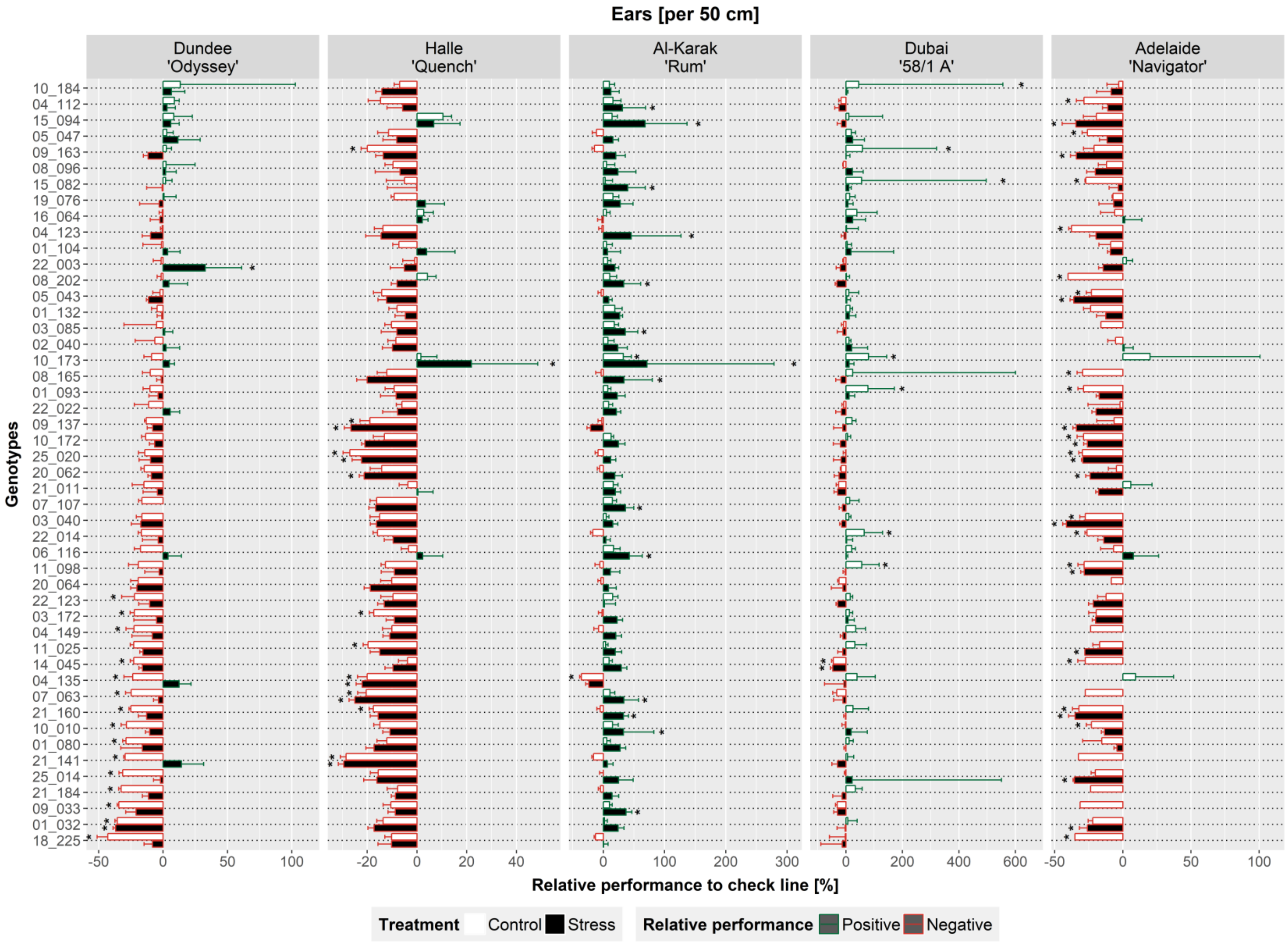

**Figure S10.** Location names and the respective check cultivar are indicated in the grey rectangles above each plot. On the x-axis, the relative performance (in %) of HEB-YIELD lines compared to the check cultivar is depicted. On the y-axis, the 48 HEB-YIELD lines are listed, sorted based on performance under control treatment in Dundee. The increase and decrease in performance relative to the check cultivar is indicated in red and green bars, respectively. Empty and filled bars indicate relative performance under control and stress treatments, respectively. Based on a Dunnett test, HEB-YIELD lines significantly (P < 0.05) deviating from the check cultivar are marked with black asterisks above the bars.

### Supplementary Figure S11. Relative grain number per ear performance of 48 HEB YIELD lines compared to a local check cultivar under control and stress treatments

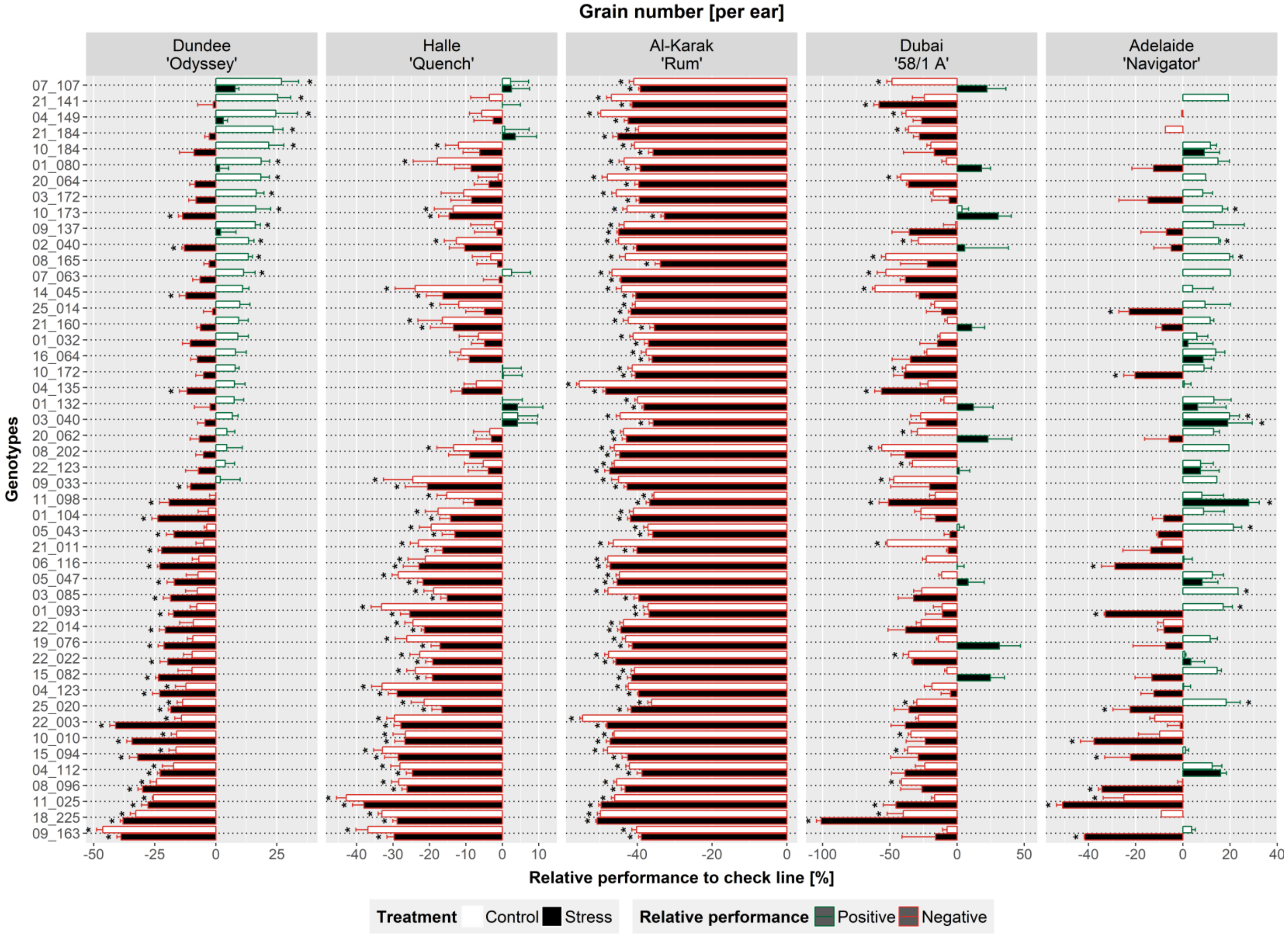

**Figure S11.** Location names and the respective check cultivar are indicated in the grey rectangles above each plot. On the x-axis, the relative performance (in %) of HEB-YIELD lines compared to the check cultivar is depicted. On the y-axis, the 48 HEB-YIELD lines are listed, sorted based on performance under control treatment in Dundee. The increase and decrease in performance relative to the check cultivar is indicated in red and green bars, respectively. Empty and filled bars indicate relative performance under control and stress treatments, respectively. Based on a Dunnett test, HEB-YIELD lines significantly ($P < 0.05$) deviating from the check cultivar are marked with black asterisks above the bars.

#### Supplementary Figure S12. Relative thousand grain weight performance of 48 HEB YIELD lines compared to a local check cultivar under control and stress treatments

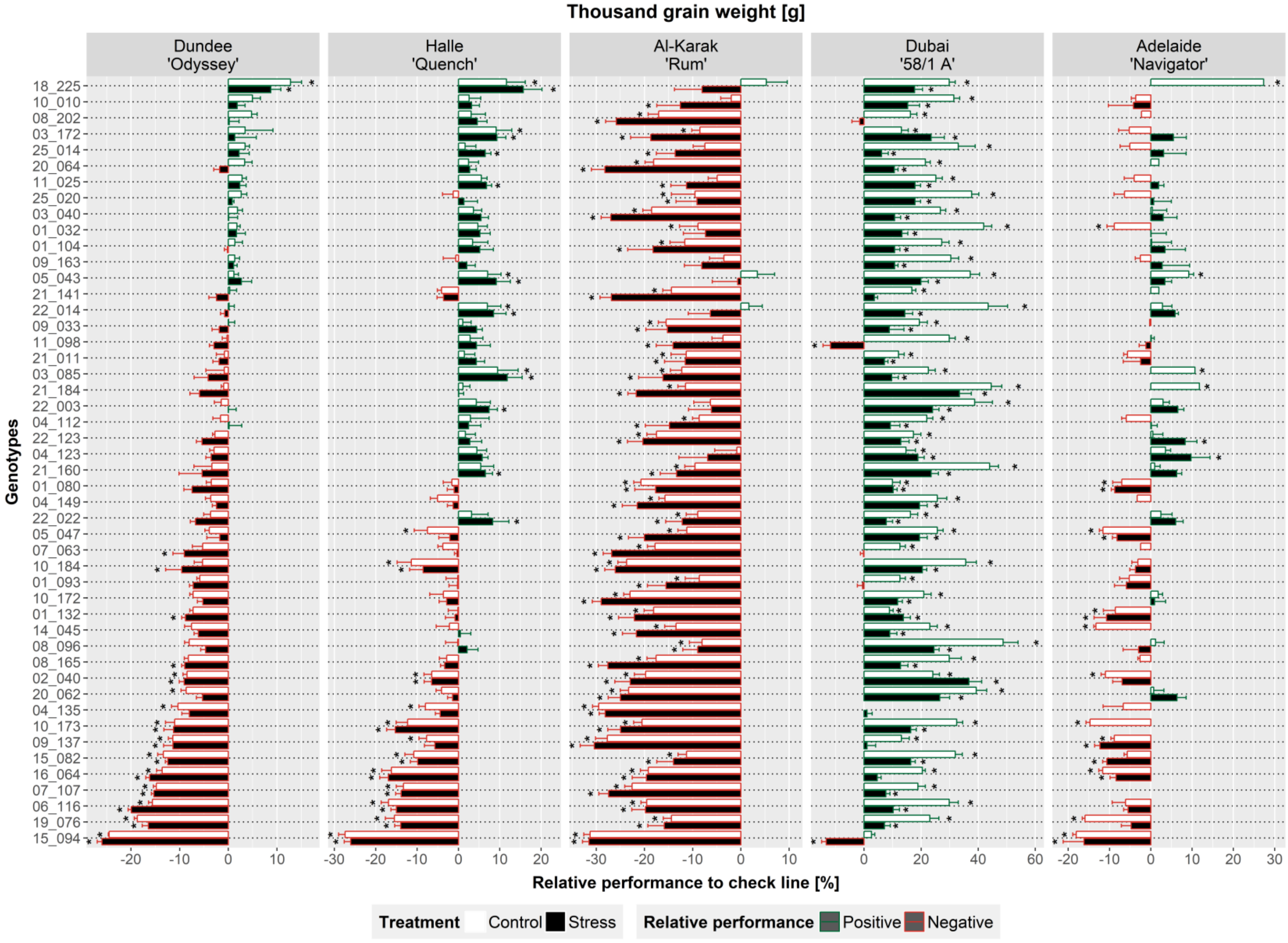

**Figure S12.** Location names and the respective check cultivar are indicated in the grey rectangles above each plot. On the x-axis, the relative performance (in %) of HEB-YIELD lines compared to the check cultivar is depicted. On the y-axis, the 48 HEB-YIELD lines are listed, sorted based on performance under control treatment in Dundee. The increase and decrease in performance relative to the check cultivar is indicated in red and green bars, respectively. Empty and filled bars indicate relative performance under control and stress treatments, respectively. Based on a Dunnett test, HEB-YIELD lines significantly ($P < 0.05$) deviating from the check cultivar are marked with black asterisks above the bars.
